# Thermotolerant pollen tube growth is controlled by RALF signaling

**DOI:** 10.1101/2025.10.25.684177

**Authors:** Rasha Althiab Almasaud, Sorel V. Yimga Ouonkap, Jared Ingram, Yahir Oseguera, Mohamed Alkassem Alosman, Cassandra Travis, Atticus Henry, Michelle Medina, Nathalie Oulhen, Gary Wessel, Alison DeLong, James Pease, Nicholas DaSilva, Mark A. Johnson

**Affiliations:** Department of Molecular Biology, Cell Biology, and Biochemistry; Brown University; 185 Meeting Street, Providence, RI 02912, USA; Department of Evolution, Ecology and Organismal Biology; The Ohio State University; Aronoff Laboratory, 318 W. 12th Ave, Columbus, OH 43210, USA

**Author notes:** **Corresponding authors:** M.A. Johnson (Lead Contact). co-first authors, contributing equally (Rasha Althiab Almasaud, Sorel V. Yimga Ouonkap).

**Keywords:** crop productivity, heat resilience, thermotolerance, signal transduction, pollen tube integrity, RALF signaling

## Abstract

Reproductive success at high temperature (HT) depends on the ability of pollen to germinate a pollen tube that delivers sperm for double fertilization. We recently found that *Solanum lycopersicum* (tomato) cultivars capable of setting fruit at HT produce pollen tubes that continue to grow under heat stress. Our present goal is to define the molecular basis of thermotolerant pollen tube growth using quantitative live imaging, transcriptomic and proteomic profiling, and genetic analysis. We found that pollen tube integrity - the ability to maintain an intact tip during germination and elongation - was the pollen performance parameter that distinguished thermotolerant cultivars. We identified the components of the Rapid Alkalinization Factor (RALF) pollen tube integrity signal transduction pathway in tomato and found that RALF peptides repress germination at control temperature and that signaling is antagonized by HT. Loss of a single RALF peptide changes pollen tube cell wall pectin distribution and enhances pollen tube integrity at HT in thermotolerant Heinz, revealing that RALF signaling regulates thermotolerant pollen tube growth and establishing a molecular framework for improving reproductive heat-resilience in crop plants.

## Introduction

Erratic growing-season temperatures cause yield loss in crops^1^, a problem expected to worsen as global food demand rises^2,3,4^. Many staple crops (wheat, rice, corn, fruits) depend on fertilization, yet this process occurs within a narrow thermal window^5–13^. Fertilization requires rapid (∼1–12 h), long (∼1000x initial diameter), highly polarized, and temperature-sensitive^14^ pollen tube growth from stigma to ovules within an ovary^15^. Pollen grains, which develop in anthers, germinate on the stigma, extend tubes guided through floral tissue^16,17^, and rupture inside ovules to release two sperm for double fertilization^18^, producing embryo and endosperm, the core components of seeds. Because seed and fruit production depend on pollen performance—the ability to germinate, elongate rapidly, and avoid premature rupture—understanding the molecular basis of thermotolerant pollen performance is critical for improving crop resilience.

Tomato is an excellent model to define how pollen performance is affected by HT because each flower produces pollen quantities sufficient for genome-scale analyses^19^, pollen tubes germinate rapidly and synchronously in vitro^20^, and cultivated tomato and its wild relatives have adapted to a wide range of temperatures^21^. We recently found that heat stress applied only during the pollen tube growth phase is sufficient to limit tomato fruit and seed production^19^. We also showed that thermotolerant cultivars, which maintain fruit and seed set at HT, have pollen tubes that grow better under HT (in the pistil and in vitro) than the Heinz reference cultivar^19^. However, these studies did not examine the early dynamics of pollen germination, tube extension, or the maintenance of pollen tube integrity—critical components of pollen performance.

Pollen tube integrity^22^ is the ability of the pollen tube to continue extending at its tip without bursting until it successfully reaches an ovule - it is essential for fertilization. Integrity is controlled by an autocrine signaling pathway in which rapid alkalinization factor (RALF) proteins^23–26^ secreted by the pollen tube are perceived on the plasma membrane by *Catharanthus roseus* RLK1-like receptors (e.g. ANXUR, ANX^22^ and Buddha’s Paper Seal, BUPS^25,27,28^) in concert with LORELEI-like glycosylphosphatidylinositol (GPI)-anchored proteins (LLG) co-receptors^27^. RALFs also directly modulate the cell wall through interactions with leucine-rich repeat extensin proteins (LRXs^26,29–31^). These signaling pathways have predominantly been studied in Arabidopsis, but RALFs were recently shown to control pollen tube integrity in Maize^32^. Here, we test whether maintaining integrity is a critical component of reproductive thermotolerance and whether RALFs contribute to pollen thermotolerance using tomato as a model.

## Results

### Pollen tube integrity distinguishes thermotolerant from thermosensitive pollen tube growth

Using live-imaging^20^, we quantified pollen performance over two hours at 28° or 34° (Fig. 1, all temperatures reported in degrees Celsius). Previously, we examined heat stress on pollen tubes after 6 hours of growth in vitro^19^; here, our goal was to analyze pollen function during the critical phases of germination and early elongation. Pollen from greenhouse-grown plants was incubated in growth media, and each grain in the field of view was classified in every frame (Fig. 1a–b). Hydrated grains (Fig. 1a,b, red square) were scored for “activation”, either tube-producing (green square) or rupturing (blue square). Extending pollen tubes were further classified as intact (green squares) or ruptured (cyan squares). Pollen tube tips were tracked across the time course to calculate extension rates (Supplementary Fig. 1). Dozens of grains were analyzed in each replicate, with at least four replicates per temperature for each of six cultivars (Supplementary Figs. 1,2). Representative results from Heinz (reference), Tamaulipas (thermotolerant), and Nagcarlang (thermotolerant) illustrate the parameters distinguishing thermotolerant cultivars^19^ (Fig. 1).

**Figure 1:**
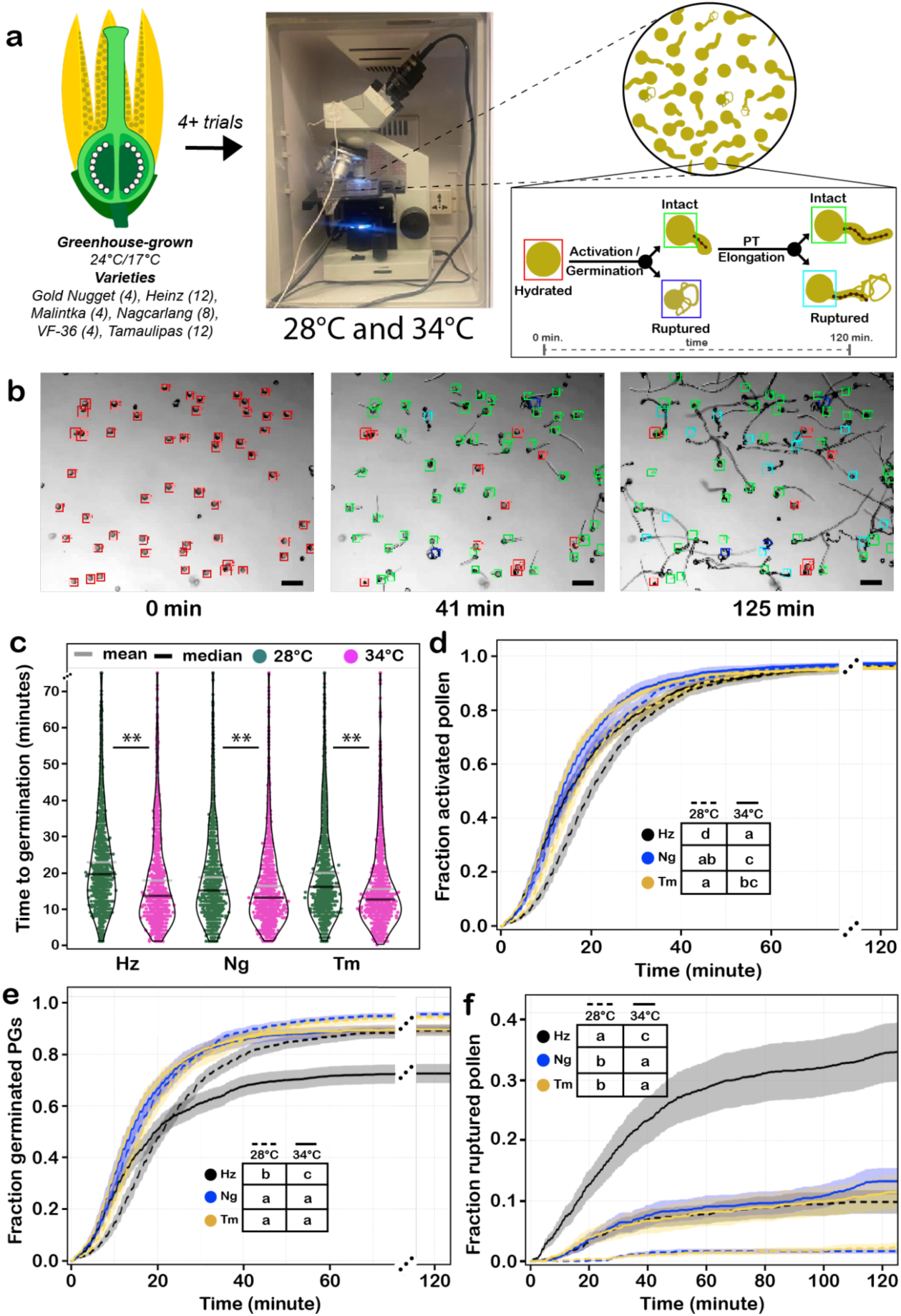
Live cell imaging of pollen growth reveals that high temperature promotes activation/germination and rupture, leading to decreased performance for thermosensitive Heinz. (**a**) Illustration of our approach to study the impact of temperature stress on pollen performance using live imaging. The numbers in parentheses (left) represent the number of trials per cultivar. Colored squares (right) represent pollen states tracked in each recording. (**b**) Representative images (Heinz) of the tracking results of germination and viability during germination and early elongation. The colored boxes indicate the state of each pollen as in (a). (**c**) Time to successful germination at indicated temperatures. Each dot represents a germinated pollen grain. Statistical analysis was performed using the Mann-Whitney-Wilcoxon U test using p.value thresholds of ns > 0.01 > * > 0.001 > **. (**d**) Fraction of activated pollen grains over time – grains that germinated a tube (green) or burst (blue or cyan). (**e**) Fraction of successfully germinated pollen grains over time (green). (f) Fraction of ruptured pollen grains and tubes over time (blue or cyan). For **d-f**, Statistical analysis was performed using survival analysis (methods) with a time cutoff of 125 minutes and pairwise p.values were bonferroni adjusted. Similar letters in tables represent no statistical significance between groups at a p.value < 0.01. Different letters in tables represent statistically significant differences at a p.value < 0.01 between groups.

We found that germination occurred significantly earlier at 34° in all cultivars (Fig. 1c), indicating HT accelerates germination. Analysis of the fraction of activated (germinated or ruptured pollen) showed a small, but significant effect of HT, with time to 50% activation faster at 34° (solid lines) than at 28° (dotted lines, Fig. 1d). However, when examining successful germination, HT significantly affected only Heinz (Fig. 1e) and VF36 (Supplementary Fig. 2c), both reaching a maximum of 65%. When we analyzed the fraction of ruptured pollen over time, we found that germination was reduced because over 30% of Heinz pollen ruptured at HT during the time course (Fig. 1f). Interestingly, this analysis also highlights that at both 28° and 34°, Tamaulipas and Nagarlang have very low pollen rupture rates that are significantly lower than Heinz. These data suggest that thermotolerant cultivars have enhanced pollen performance at HT because they maintain pollen tube integrity during activation and elongation.

### Thermotolerant Nagcarlang modulates its pollen transcriptome and proteome in response to germination, but is relatively unaffected by HT

When we previously analyzed HT-induced gene expression in pollen tubes grown for six hours in vitro^19^, we found evidence of stress-response priming: thermotolerant Tamaulipas had higher basal levels of genes that were induced by HT in Heinz^19^. To test whether priming is also a feature of germination and early elongation, we collected transcriptome and proteome data over two hours (Fig. 2a; blue stars, RNA-seq; green stars, proteome). Principal component analysis showed genotype is the primary driver of variance in global mRNA and protein expression (Fig. 2b; this was also observed in pollen tubes grown for six hours^19^). Transcriptome variance increased with time under HT (mature pollen vs. 120 min, Fig. 2b) as pollen tubes extended and responded transcriptionally, but proteome variance remained limited (Fig. 2b, Proteome).

**Figure 2:**
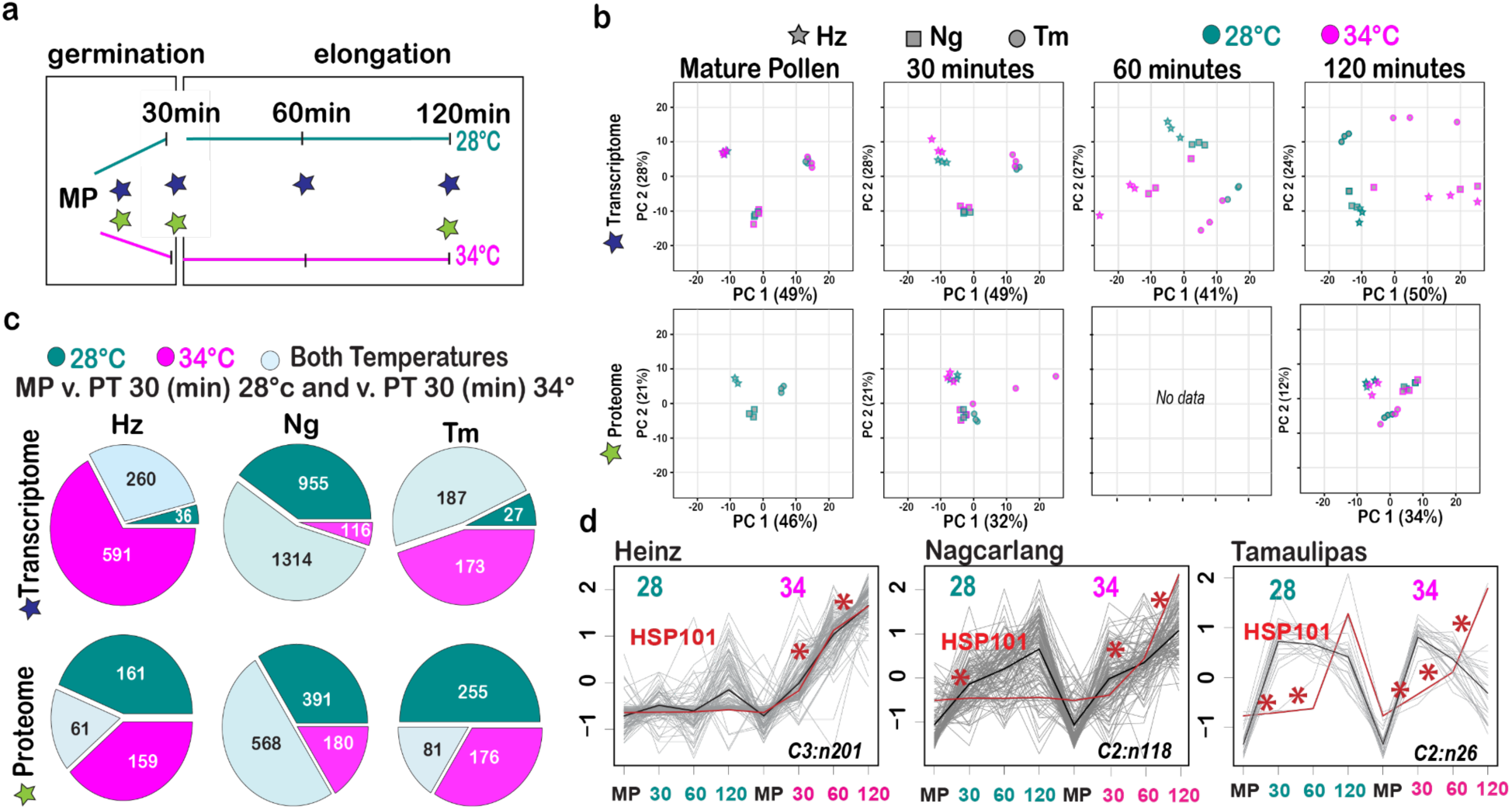
Germinating pollen of thermotolerant cultivars exhibit distinct transcriptomic and proteomic responses to HT. **(a)** Time points used for transcriptome (blue star) and proteomic (green star) analysis. **(b)** PCA was performed on transcriptomic data from mature pollen samples at 30, 60, and 120 min after germination, and on proteomic data at 30 and 120 min post-germination, across Heinz (Hz), Nagcarlang (Ng) and Tamaulipas (Tm) cultivars. Each point on the plot represents a sample, with its position determined by the first two principal components capturing the most significant variance in the data. **(c)** Pie chart presents the distribution of differentially expressed genes (DEGs) and proteins (DEPs) between mature pollen and pollen tubes at 30 min, at 28° and 34° across three cultivars. **(d)** The transcriptomic cluster (Suppl. Fig. 3) containing heat stress indicator, HSP101, for each cultivar. The cluster number and the number of genes in each cluster is provided. Red stars denote statistically significant differences in expression between time points.

Because avoiding rupture during germination is critical for thermotolerance (Fig. 1e), we compared differentially expressed genes (DEGs) and proteins (DEPs) between mature pollen and pollen that had germinated (30 minutes) at 28° or 34° (Fig. 2a). Thermotolerant Nagcarlang expressed ∼3× more germination-associated DEGs and DEPs than Heinz or Tamaulipas (Fig. 2c). Heinz showed the largest proportion of DEGs/DEPs altered only at HT, while Nagcarlang had the fewest (pink, Fig. 2c), suggesting Nagcarlang broadly activates the genome during germination, with expression less perturbed by HT.

DEGs exceeded DEPs in all cultivars (Supplementary Fig. 3a) and overlap was low (≤15%; 83/568 in Nagcarlang, Supplementary Fig. 3a). While this disconnect could be partly due to higher RNA-seq sensitivity compared to proteomics, this finding may also suggest HT-induced RNA changes were not translated into protein changes, especially in thermosensitive Heinz. We used k-means clustering to define gene groups with shared patterns during germination at both temperatures (Supplementary Figs. 4,5; Supplementary Tables 1,2) and, interestingly, stress response genes clustered differently across cultivars. HSP101, the first HSP induced (30 min, 28° vs. 34°; Supplementary Fig. 6), marked stress responses in all cultivars, but clustered differently: in Heinz, with genes induced only at HT (Fig. 2d); in Nagcarlang and Tamaulipas, with genes responsive to germination at both temperatures. These data are consistent with the proposal that optimal germination and pollen tube elongation require expression of genes that have been classically defined as stress-responsive^33,34^ and add perspective to our recent finding that the transcriptome of pollen tubes grown in vitro for six hours was primed for stress responses in thermotolerant Tamaulipas^19^. By analyzing germination and early pollen tube growth (Fig. 2) we now find that many stress-responsive genes are activated by germination in thermotolerant cultivars (Fig. 2d) and thus appeared to have been primed in our prior analysis of six hours of pollen tube growth^19^.

### Analysis of the tomato pollen tube RALF signaling pathway

Pollen tube integrity during germination and elongation distinguishes thermotolerant from thermosensitive tomato cultivars (Fig. 1f). To test whether the pollen tube integrity-controlling^25–27,32^ RALF signaling pathway underlies thermotolerant pollen tube growth in tomato (Fig. 1f), we defined pollen expressed RALFs (Fig. 3a), and their potential receptors and co-receptors (Supplementary Fig. 7). The tomato genome encodes 10 RALF genes (SlRALF1–10, Supplementary Table 3) and we constructed a phylogeny with 37 Arabidopsis RALFs^35^, to identify tomato candidates likely regulating pollen tube integrity. *SlRALFs 1-4* were abundantly expressed in mature pollen and pollen tubes and *SlRALFs 1, 3, and 4* form a small clade related to *AtRALFs 4 and 19* (Fig. 3a) - the core ligands for the autocrine signaling pathway controlling pollen tube integrity in Arabidopsis^25,26^. Notably, *SlRALF3* was previously characterized (SlPRALF) and shown to control pollen tube elongation^24^.

**Figure 3:**
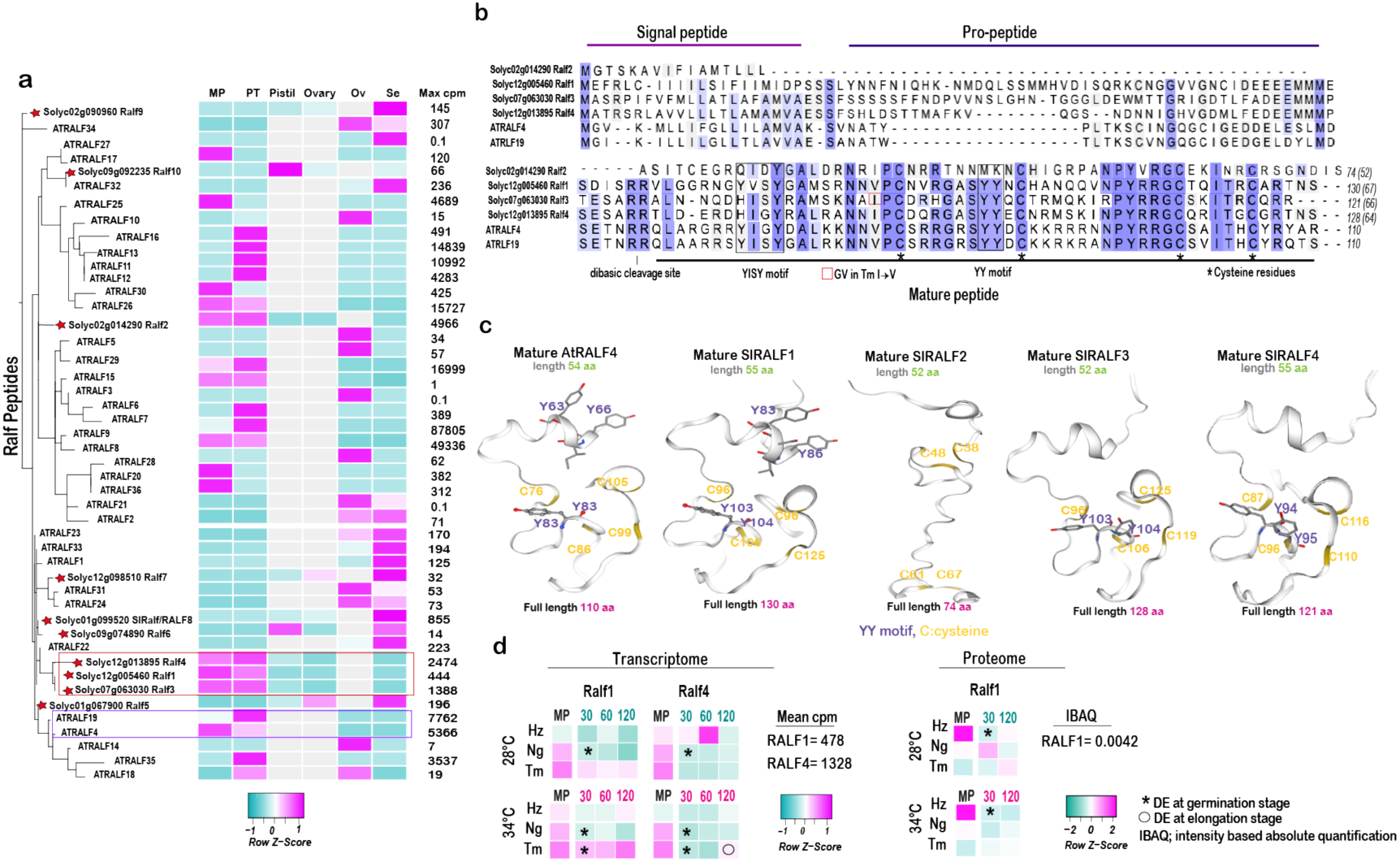
Tomato pollen expresses four RALFs. **(a)** Phylogenetic tree illustrating the evolutionary relationships between tomato peptides and Arabidopsis Ralfs peptides. Additionally, a heatmap shows the expression levels of these peptides in different tissues such as mature pollen "MP", pollen tube "PT", pistil, ovary, ovule "Ov" and seedling "Se". Max cpm represents the highest expression value observed across the different tissues. Gray boxes in the heatmap indicate tissues for which data are missing or not provided. **(b)** Peptide alignment comparing the most abundant RALF peptides found in tomato mature pollen with Arabidopsis pollen-expressed RALFs. The length of the predicted mature peptides is shown in parentheses following the total amino acid count. Alignments were visualized using the UniProt alignment tool (https://www.uniprot.org/align). **(c)** Structural models of the mature peptides of Arabidopsis RALF4 (AtRALF4) and tomato SlRALF1, SlRALF2, SlRALF3, and SlRALF4 highlight conserved YISY and YY ring motifs essential for receptor binding, along with cysteine residues (C). The models were generated using SWISS-MODEL. **(d)** Heatmap representation showcasing the expression levels of RALF1 and RALF4 (data for RALF2 and 3, Supplementary Fig. 7), at both transcriptomic and proteomic levels across various time points and cultivars, under control and heat stress conditions. The significance of expression changes is denoted by stars (*) at the germination stage and circles (o) at the tube elongation stage. Colored time point labels indicate cyan for control temperature (28°) and magenta for high temperature (34°).

SlRALF1, SlRALF3, and SlRALF4, like AtRALF4 and AtRALF19 (Fig. 3b), have sequences associated with RALF processing (a dibasic cleavage site for pro-peptide removal^23,24^), four conserved cysteine residues essential for proper folding^23,24^, and a YY (double tyrosine) motif implicated in interaction with CrRLK1L receptor kinases^36^. In contrast, SlRALF2, the RALF with the highest mRNA abundance in tomato pollen (Fig. 3a), was in a clade by itself (Fig. 3a) and lacked these characteristic motifs (Fig. 3b), suggesting potential functional divergence. Structural modeling of mature peptides further underscores significant differences in SlRALF2 compared to the other peptides (Fig. 3c). Analysis of RALF genetic variation across thermotolerant and thermosensitive cultivars^19^ identified a single amino acid variant in SlRALF3, where isoleucine (I^94^) is replaced by valine (V) in the Tamaulipas cultivar.

RNA-seq analysis of *SlRALF1-4* levels during pollen germination and elongation (Fig. 3d) indicated that RALF peptides are not induced by heat stress, but did reveal significant decreases in transcript levels during pollen germination (Fig. 3d). Only SlRALF1 was detected in our proteome analysis and it was found to significantly decrease during germination in Heinz at both temperatures (Fig. 3d). These data indicate that RALF expression levels are not affected by growth at HT, but that decreases in RALF expression could be important for pollen germination, which is in keeping with prior work showing that treatment with RALF peptides inhibits pollen tube^24,26,27,37^ and root hair growth^38^.

### SlRALF treatment represses pollen activation and is opposed by HT

To test whether tomato pollen RALFs are active and whether activity is altered by HT and/or genetic background, we synthesized mature SlRALF peptides (1-4) and analyzed their effects on Heinz, Tamaulipas, and Nagcarlang pollen growth using our live-imaging system (Fig. 4). We used 1 μM RALF peptide because end-point analysis suggested this concentration would facilitate analysis of the kinetics of pollen germination, tube growth, and integrity; higher concentrations completely block pollen germination (Supplementary Fig. 8). SlRALF2, which had no activity in endpoint assays (Supplementary Fig. 8) and no RALF treatment (Fig. 4, Supplementary Fig. 9) were used as negative controls. Pollen performance was classified over two hours at 28° or 34° (Fig. 4a).

**Figure 4:**
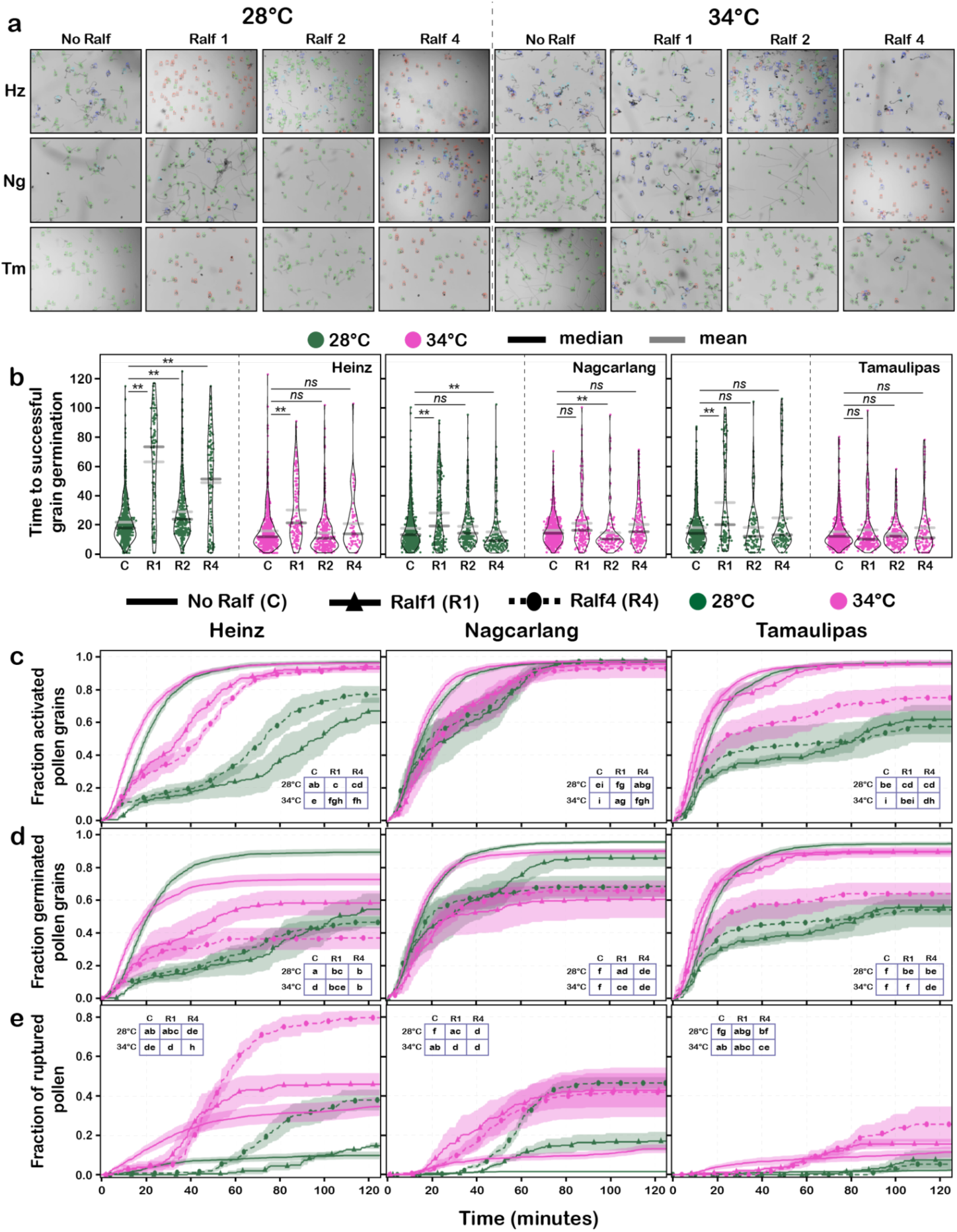
Effects of Ralfs and Temperature on pollen grain activation, germination, and cell rupture. **(a)** Representative images of pollen tube growth and data quantification after 60 minutes of growth. **(b)** Time to successful germination at indicated temperatures and in the presence of 1uM Ralf 1, Ralf 2, or Ralf 4. Each dot represents a germinated pollen grain. Statistical analysis was performed using the Mann-Whitney-Wilcoxon U test using p.value thresholds of ns > 0.01 > * > 0.001 > **. **(c)** Fraction of activated pollen grains over time – grains that exited the hydrated state (Figure 1a, red). **(d)** Fraction of successfully germinated pollen grains over time (Figure 1a, red to green). **(e)** Fraction of ruptured pollen grains and tubes over time (Figure 1a, blue + cyan). For c-e, Statistical analysis was performed using survival analysis (methods) with a time cutoff of 125 minutes and pairwise p.values were bonferroni adjusted. Similar letters in tables represent no statistical significance between groups at a p.value < 0.01. Different letters in tables represent statistically significant differences at a p.value < 0.01 between groups.

At 28°, SlRALF1 significantly delayed germination in all cultivars and SlRALF4 had a similar effect, but only on Heinz (Fig. 4b). These delays were significantly diminished at 34°, indicating that HT antagonizes RALF repression. SlRALF1 and SlRALF4 also delayed pollen activation (germinated + ruptured pollen grain), most strongly in Heinz and Tamaulipas, where activation never exceeded 60% (Fig. 4c). Nagcarlang pollen, by contrast, was more resistant to RALFs: 50% activated within 40 minutes under SlRALF1 treatment, compared with 100 minutes for Heinz. At 34°, activation accelerated for treated samples in all cultivars, again showing that HT counteracts RALF repression.

Analysis of successful germination confirmed these trends: about half of SlRALF1- or SlRALF4-treated Heinz pollen failed to germinate in two hours, whereas Nagcarlang was significantly less repressed (Fig. 4d). Analysis of rupture kinetics revealed a two-phase response: during the first 35 minutes of germination, SlRALF4 suppressed rupture at both temperatures, but afterward rupture increased in treated samples (Fig. 4e, dotted lines with circles). In Heinz, SlRALF4 promoted rupture over time at both 28° and 34°, with 80% ruptured by 90 minutes at HT (Fig. 4e). SlRALF1 (solid lines with triangles) had little effect on rupture. In Nagcarlang, SlRALF4 promoted rupture at both temperatures, but effects did not synergize with HT as they did in Heinz. In thermotolerant Tamaulipas, rupture remained minimal, with only mild increases under RALF treatment (Fig. 4e).

Together, these results show that RALF1 and RALF4 repress germination and activation, effects that are antagonized by HT and modulated by cultivar, with Nagcarlang showing the greatest resistance. We propose that pollen tube RALF signaling is diminished by HT and that thermotolerant cultivars have optimized RALF signaling to maintain pollen tube integrity at HT.

### Loss of SlRALF1 increases pollen tube integrity at HT in Heinz and delays germination in Tamaulipas

To test whether RALF signaling regulates thermotolerant pollen tube integrity, we generated *SlRALF1* loss-of-function mutants in Heinz (*ralf1-6^H^*) and Tamaulipas (*ralf1-4^T^*, Fig. 5a). We focused on SlRALF1 because it was detectable in our pollen proteomic analysis (Fig. 3e) and had peptide treatment effects distinct from those of SlRALFs 2-4 (Fig. 4). Prior studies in Arabidopsis^25,26^ and Maize^32^ suggested that loss/reduction of two pollen *RALFs* was necessary to reveal mutant phenotypes, so we tested whether quantitative live-imaging under HT in tomato could detect individual *RALF* functions.

**Figure 5.**
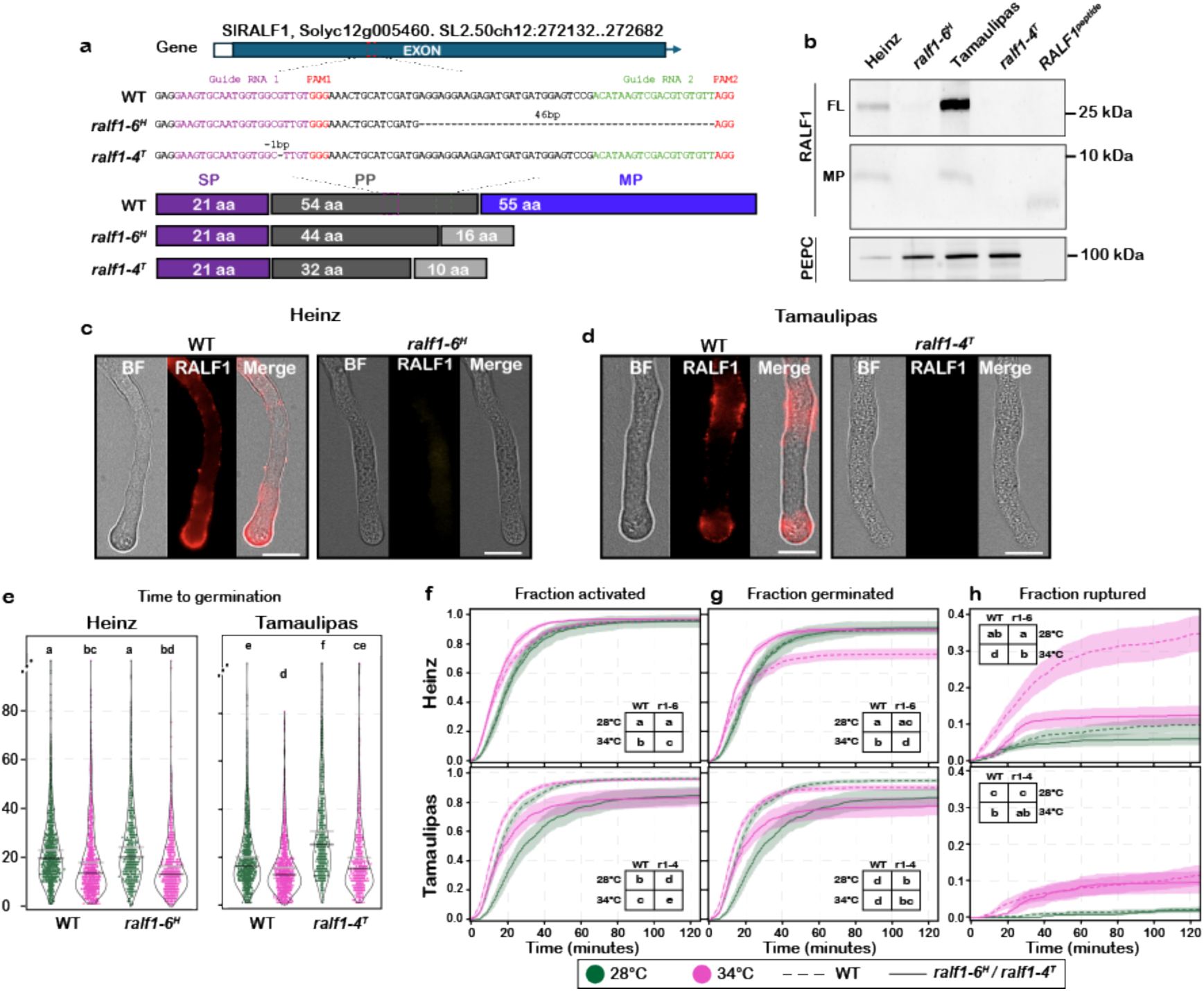
RALF1 localization and pectin deposition in tomato pollen tubes. **(a)** Schematic of CRISPR-Cas9 generated *ralf1* mutants in Heinz (*ralf1-6^H^*) and Tamaulipas (*ralf1-4^T^*) showing deletion of the mature peptide coding sequence. **(b)** Western blot analysis was performed using anti-RALF1 antibodies on protein extracts from pollen tubes grown at 28°C of WT Heinz, WT Tamaulipas, *ralf1* mutant lines, and synthetic RALF1 peptide. Phosphoenolpyruvate carboxylase (PEPC) was used as a loading control for the same samples. FL: full-length and MP: mature peptide. **(c, d)** Representative immunolocalization images of RALF1 at 90 minutes after pollen germination in Heinz and Tamaulipas, respectively. **(e)**, fraction of activated pollen **(f)**, germinated pollen **(g)**, and ruptured pollen **(h)** over time. Statistical significance was determined using the Kruskal–Wallis test and Man-Withney-Wilcoxson (e) and survival analysis (f-h) and resulting pairwise p.values were Bonferroni adjusted. For each panel (e-h), similar letters indicate no statistical significance at p < 0.01 and different letters indicate statistical significance at p < 0.01.

Immunoblots with anti-RALF1 antibodies detected precursor and mature peptides in wild-type samples; both were absent in mutants (Fig. 5b, Supplementary Fig. 10a,b). Immunolocalization showed that SlRALF1 accumulation in the shank and tip of wild-type pollen tubes was not detectable in mutants (Fig. 5c-d).

Live-imaging at 28° and 34° revealed cultivar- and temperature-specific mutant phenotypes (Fig. 5e–h). In Heinz, *ralf1-6^H^* had little effect at 28° but enhanced pollen tube integrity at 34° - pollen rupture plateaued at ∼10% by 30 minutes at 34° in *ralf1-6^H^* (similar to Tamaulipas at 34°), but reached ∼35% by 2 hours in wild-type (Fig. 5h). Successful germination at 34° also improved, reaching ∼85% in *ralf1-6^H^* mutants versus ∼70% in wild-type (Fig. 5g). Thus, loss of *SlRALF1* improves Heinz pollen tube integrity at HT.

In Tamaulipas, *ralf1-4^T^* did not affect rupture (Fig. 5h) but significantly slowed activation (Fig. 5f) and germination (Fig. 5g) at both temperatures. Mutant pollen tubes also germinated significantly later than wild-type at both temperatures (Fig. 5e). Therefore, *SlRALF1* reduces thermotolerance in Heinz by compromising pollen tube integrity under HT, but in thermotolerant Tamaulipas it modulates the kinetics of germination across a range of temperatures.

### RALF-regulated pollen tube pectin distribution is altered by growth at HT

Having established that HT disrupts pollen performance by decreasing pollen tube integrity (Fig. 1), that the effects of RALF treatments on pollen tube growth are counteracted by HT (Fig. 4), and that loss-of-function of *SlRALF1* enhances thermotolerant pollen tube growth in Heinz (Fig. 5), we set out to address how HT and RALF signaling interact to control deposition of pectin, a key component of the pollen tube cell wall^39,40^. Methyl-esterified pectin is relatively flexible and enriched at the extending pollen tube tip; whereas de-esterified pectin is relatively rigid and is enriched in the pollen tube shank^39,41^. Recently, it was shown that maize pollen tube RALFs are required for proper distribution of pectin to maintain pollen tube integrity^32^. We analyzed pectin distribution using LM19 (de-esterified pectin-specific antibody^42^) and LM20 (methyl-esterified pectin-specific antibody^42^) immunolocalization in pollen tubes grown in vitro for 2 hours at 28° or 34° (Fig. 6a).

**Figure 6:**
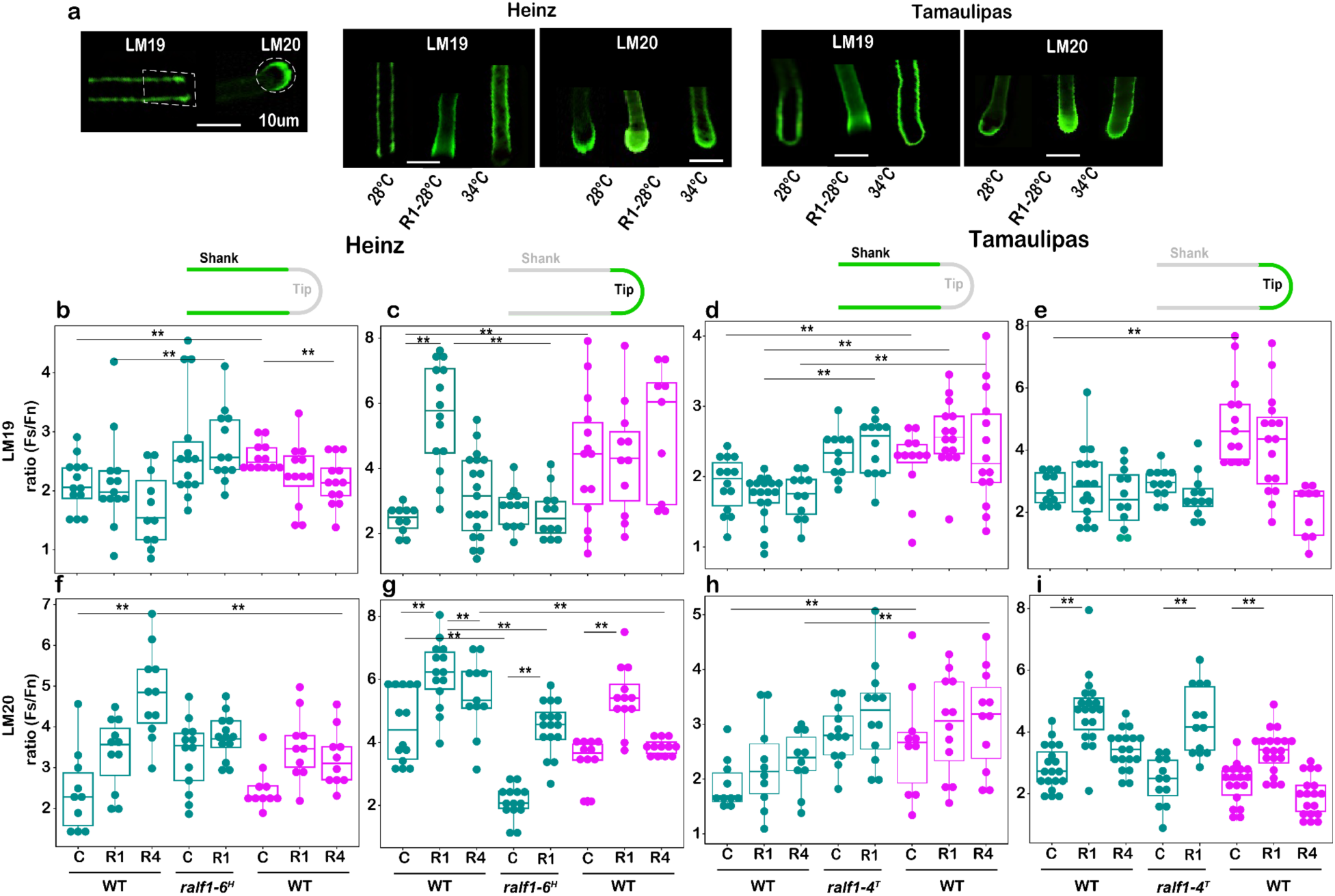
RALF signalling and temperature interact to control pectin localization in pollen tubes. **(a)** Representative images of LM19 and LM20 immunostaining, showing pectin distribution in the shank and tip regions of pollen tubes, respectively, across tomato cultivars. RALF1 (R1) treatment is used as a representative for staining patterns. Dashed lines indicate the regions of interest (ROI) measured for fluorescence intensity. **(b-i)** Quantification of fluorescence intensity for LM19 and LM20 staining in the shank and tip region under 1µM RALF1 (R1), or 1µM RALF4 (R4) treatments wild-type (WT) or mutant Heinz and Taumalipas at 28° (green) and 34° (magenta). Fluorescence intensity was normalized using the ratio of LM19 or LM20-stained fluorescence (Fs) to fluorescence of unstained controls (Fn). Statistical significance was determined using Dunn test and indicated as p < 0.01 (**), and ns (not significant).

First, we addressed how temperature affects the distribution of pectin in growing pollen tubes, comparing patterns in thermosensitive Heinz and thermotolerant Tamaulipas. LM19 fluorescence intensity increased significantly in the shank and tip under HT in Heinz and Tamaulipas (Fig. 6b-e; green, 28°; magenta, 34°), indicating an increase of de-esterified (rigid) pectin across the pollen tube. LM20 fluorescence is highest at the tip and levels of tip-focused LM20 signal were not significantly changed by HT in Heinz or Tamaulipas (Fig. 6a,g,i). However, LM20 levels in the shank significantly increased with HT in thermotolerant Tamaulipas (Fig. 6h). These data, along with similar analysis of Nagcarlang (Supplementary Fig. 10c,d), indicate that HT changes pectin distribution in the pollen tube cell wall and that these changes vary across thermotolerant and thermosensitive genotypes.

Next we addressed how RALF treatment affects pectin distribution. Treatment with 1µm SlRALF1 significantly increased the levels of LM20 in the tip of Heinz and Tamaulipas when grown at either 28° or 34° (Fig. 6g,i). This was not observed for either SlRALF4 (Fig. 6g,i) or SlRAFL2 (Supplementary Fig. 10c), suggesting a specific function for SlRALF1. Moreover, in *ralf1-6^H^* mutant pollen tubes, we found that LM20 levels significantly decreased in the tip relative to wild-type Heinz and that mutant levels could be restored by addition of exogenous SlRALF1 to mutants (Fig. 6g). Treatment with 1uM SlRALF1, SlRALF4 (Fig. 6b,d) or SlRALF2 (Supplementary Fig. 10d) did not significantly alter the LM19 signal in the shank of pollen tubes grown at 28° in any cultivar and loss-of-function of SlRALF1 (*ralf1-6^H^* or *ralf1-4^T^)* did not affect the LM19 signal in the shank, relative to wild-type controls at 28° (Fig. 6b,d). These data indicate that RALF1 is specifically required to regulate levels of methyl-esterified pectin at the pollen tube tip.

## Discussion

We recently found that HT applied only during the pollen tube growth phase was sufficient to reduce fruit and seed production in thermosensitive cultivars of tomato, but that thermotolerant cultivars had better pollen function at HT^19^. Here, we’ve shown that the ability to maintain pollen tube integrity at HT is the pollen performance parameter that best distinguishes thermotolerant from thermosensitive cultivars (Fig. 1, 7) and that RALF signaling controls thermotolerant pollen tube integrity (Figs. 4 and 5, 7). Thus, we establish a mechanistic framework for heat-resilient reproduction in flowering plants and provide an avenue to develop crop varieties that can maintain yield when heat waves occur during the pollination season.

### Thermotolerant signal transduction is required for resilient cellular function in plants

Plant cells have evolved sophisticated mechanisms to sense and respond to changing temperature, light, and water availability throughout each day - and extreme variation of these conditions throughout the growing season^43,44^. The pollen cell is a useful model to understand molecular responses at the cellular level because we can measure many parameters of pollen cellular performance (Fig. 1, Supplementary Figs. 1,2) under control and HT. We found that loss of *SlRALF1* in thermosensitive Heinz increased pollen performance at HT by enhancing pollen tube integrity (Fig. 5h) but had modest effects on thermotolerant Tamaulipas, and that HT and RALF signalling are antagonistic (Fig. 4, 5). These data suggest that the RALF signalling outputs change across a range of temperatures. One possibility is that HT reduces RALF signaling by altering the interactions among the core components of the plasma-membrane localized signaling complex (Fig. 7). Under this model, thermotolerance could result from genetic variation in signaling components that strengthen these molecular interactions at HT. We’ve observed focused genetic variation among RALFs (Fig. 3b) and receptors (Supplementary Fig. 7d) in the small set of tomato cultivars we’ve analyzed and future work will be aimed at testing whether this variation modulates thermotolerant pollen tube integrity. We can also analyze downstream components of the RALF signaling pathway to explore the basis for thermotolerant pollen tube growth. For example, we recently found that Tamaulipas pollen tubes maintain reactive oxygen species (ROS) at optimal levels under HT^19^. It has been shown that RALF signaling regulates ROS by modulating the activity of Respiratory Burst Oxidase Homolog (RBOH) enzymes ^45,46^, it will be interesting to test whether this enhanced ROS homeostasis in Tamaulipas is mediated RALF-dependent ROS regulation and/or by enhanced production of antioxidants ^47,48^.

**Figure 7.**
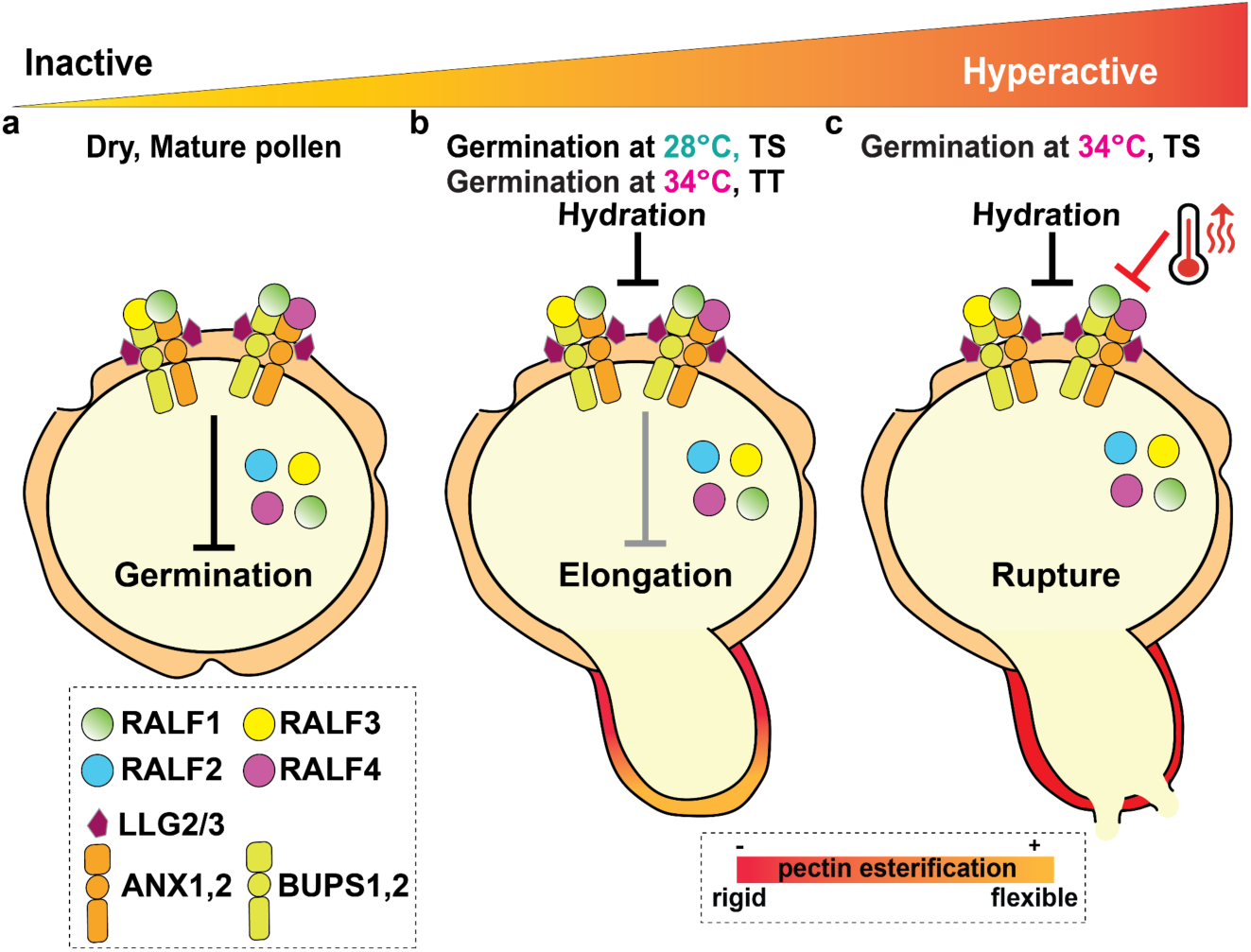
RALF signaling controls thermotolerant pollen tube growth. **(a)** In mature pollen, high levels of pollen-expressed RALF1, RALF3 and RALF4 bind their receptors ANX1/2 and BUPS1/2 and the co-receptor LLG2/3, inhibiting pollen germination (Inactive). **(b)** Upon pollen hydration, levels of RALF1 decrease, reducing inhibition and allowing pollen tube elongation. A tightly regulated balance of RALF peptides is crucial for proper germination; once the pollen tube emerges, RALF1 regulates tip-localized pectin deposition to maintain pollen tube integrity. RALF signaling has been tuned to maintain pollen tube integrity at HT in thermotolerant (TT) genotypes. **(c)** Under high temperature (34°), RALF signaling is disrupted in thermosensitive (TS) genotypes, leading to rupture (hyperactivation); but is maintained in thermotolerant cultivars. This differential thermoresponse is associated with variations in cell wall composition and pectin distribution between cultivars. The schematic illustrates dynamic changes in the distribution of esterified and de-esterified pectin across the pollen tubes that is required to maintain pollen tube integrity across a range of temperatures.

### HT and RALF signaling have opposing effects on pollen activation

We propose that pollen hydration, by incubation in growth media in vitro (Fig. 1) or by deposition on the wet tomato stigma^49^, leads to rapid RALF-controlled activation of pollen to facilitate germination and extension of the pollen tube without rupture (Fig. 7). In Arabidopsis, which has a dry stigma, RALF23/33 signaling from the stigma regulates the hydration process by activating FER-dependent, stigma-expressed, RBOH pathways to restrict water transfer to pollen during pollen–stigma interactions ^50^. We propose this dynamic could be different on the tomato wet stigma where water for hydration is freely available. Quantitative live imaging revealed that HT accelerated pollen activation (successful germination or rupture of a hydrated pollen grain) by decreasing the time to germination (Fig. 1c) and by increasing the number of pollen grains that burst in thermosensitive Heinz (Fig. 1d-f). In contrast, thermotolerant Tamaulipas maintained a larger fraction of pollen grains that successfully germinated tubes, and had a significantly higher proportion of tubes that maintained pollen tube integrity under HT (Fig. 1f). This analysis showed that maintaining integrity during germination is critical for thermotolerance. Treatment with exogenous SlRALF1, 3, or 4 suppressed pollen activation at 28° (Fig. 4c, Supplementary Fig. 8). This is an important observation suggesting that the function of autocrine RALF signaling is to suppress activation (Fig. 7) and thus maintain integrity during germination. Importantly, HT counteracted this effect of exogenous RALF treatment suggesting that HT reduces the ability of RALF signaling to properly modulate pollen activation (Fig. 7c). In the future, we will explore whether RALF signaling in the cells of the tomato stigma and transmitting tissue contribute to regulation of successful pollen tube germination and elongation.

### SlRALF1, 3, and 4 have distinct functions in controlling thermotolerant pollen tube integrity

Analysis of the Arabidopsis model system established an autocrine signaling pathway mediated by pollen-expressed AtRALF4 and AtRALF19 that contoIs pollen tube integrity^22,25–27^. *ralf4, ralf19* double mutant pollen tubes rupture prematurely in vitro and fail to grow in the pistil^25,26^. Arabidopsis pollen tubes express 12 additional RALF proteins^51^ along with two co-receptors (LLG2 and LLG3^27,52^) and four CrRLK1L receptor kinases (ANX1, ANX2, BUPS1, BUPS2^25,27^) - an important challenge is to understand the distinct functions governed by this large array of signaling proteins - progress has been made by studying closely related groups of genes using higher-order reverse-genetic analysis^51,53^. Tomato pollen expresses four RALFs, a single co-receptor (SlLLG2/3) and four CrRLK1L receptor kinases (Fig. 3, Supplementary Fig. 7) and we have begun to define functions for individual RALF proteins using quantitative analysis of pollen tube growth, heat stress, application of exogenous peptides, loss-of-function analysis, and analysis of patterns of pectin distribution.

We found that treatment with SlRALF1 or SlRALF4 peptides inhibited germination across cultivars, but also noted that responses varied, suggesting genetic variation in pollen tube RALF signaling within Tomato (Figs. 4, 7b). Interestingly, RALF4 treatment resulted in a significantly higher fraction of ruptured pollen tubes beginning at 60 minutes of pollen tube growth (Fig. 4e). This trend was observed in all cultivars and at control and HT and was not observed with RALF1 treatment (Fig. 4e). These data suggest that individual pollen tube-expressed RALFs have specific roles in regulating pollen tube integrity. In keeping with this idea, we observed distinct cellular phenotypes due to loss-of-function of a single RALF. In Heinz, loss of *SlRALF1* (*ralf1-6^H^*) enhanced pollen tube integrity at HT (Fig. 5h, fraction ruptured is significantly reduced in the mutant at HT), effectively phenocopying thermotolerant Tamaulipas. In contrast, loss of *SlRALF1* in Tamaulipas delayed germination (Fig. 5g). These data identify cultivar-specific roles for *SlRALF1* and suggest RALF signaling is optimized for germination in thermotolerant Tamaulipas, but loss of SlRALF1 in Heinz may alter RALF signaling output such that pollen tube integrity is increased under high temperature.

In growing pollen tubes, dynamic distribution of esterified and de-esterified pectin defines a mechanical gradient required to maintain pollen tube integrity as the tip rapidly extends^39,54,55^ (Fig. 7b). Our immunolabelling experiments demonstrate that HT alters this balance, increasing de-esterified (rigid) pectin in the tip and shank and enhancing esterified (flexible) pectin in the shank of thermotolerant Tamaulipas (Figs. 6, 7c). We also found that treatment with exogenous RALF peptides had distinct effects on pectin distribution. Treatment with SlRALF1 but not SlRALF4 significantly increased esterified pectin at the tip in Heinz and Tamaulipas (LM20, Fig. 6g,i). However, loss of *SlRALF1* only resulted in reduced levels of esterified pectin in the tip of Heinz *ralf1-6^H^* mutants (Fig. 6g), but not Tamaulipas *ralf1-4^T^* mutants (Fig. 6g). This result is interesting because it supports our proposal that RALF signaling has been tuned to achieve thermotolerant pollen tube growth in thermotolerant cultivars. Analysis of additional RALF single mutants and higher-order mutants in thermotolerant and thermosensitive cultivars will define how each pollen tube RALF contributes to thermotolerant pollen tube integrity.

## Materials and Methods

### Plant materials and growth conditions

*Solanum lycopersicum* (tomato) plants of the Heinz cultivar were grown under a 13-h-day/11-h-night cycle in the Brown University Research Greenhouse. Plants were propagated either from rooted cuttings of self-fertilized lines or from seeds. Seeds of Heinz 1706-BG (LA4345), Tamaulipas (LA1994), and Nagcarlang (LA2661) were obtained from the Tomato Genetics Resource Center, UC Davis. For seed-based propagation, tomato seedlings were germinated in potting soil under standard greenhouse conditions (16-h-day/8-h-night cycle, 25 °/18 ° day/night temperature) in a controlled growth chamber for two weeks before being transferred to two-gallon pots. Plants were fertilized with Israel Chemicals Limited Peters Professional 5-11-26 Hydroponic Special and adequately watered. Daily temperatures were weather-dependent, fluctuating between 21° and 28° during the day and 16.5° to 18° at night, but were relatively regulated to optimal growth conditions. To maximize flower production, fruits and stems were regularly trimmed.

### Live imaging of pollen tube performance in vitro

Less than 30 minutes prior to live imaging, pollen was collected from 4 to 5 flowers using a modified vibrating toothbrush, added to a pre-warmed pollen growth medium, place in a secondary contained filled with temperature ready water and placed under a microscope inside a temperature ready incubator for imaging for 125 minutes as explained in Ouonkap et. al. 2025^20^. Movie recordings were performed at 14 frames per second then reduced to 125 frames 1 minute apart from one another before quantification with TubeTracker^20^ with default settings. All automated qualifications were manually checked to correct for mistakes made through automated steps. Once pollen in each frame was classified as hydrated (red), ruptured as a grain (blue), ruptured as a tube (cyan) and viable and going (green, Fig. 1a,^20^) and raw files containing the information about each grain were then integrated together in R to extract information about pollen tube activation, germination, tip elongation and pollen cell integrity over time. All raw files, videos, and analysis scripts are available in supplementary data and TubeTracker is available on github at https://github.com/souonkap/TubeTracker.

### RNA extraction and RNAseq analysis

Dry mature grains and pollen tube RNA were extracted as follows: Pollen was collected from young flowers and grown in vitro according to the method described in Ouonkap et al. 2024^19^. The collected pollen was then transferred to 2 mL Eppendorf tubes by centrifugation and immediately flash frozen in liquid nitrogen. Pollen tube samples were ground using a TissueLyser II (Qiagen) for 1 min. RNA isolation was performed using the ReliaPrep™ RNA Tissue Miniprep System kit (Promega). Subsequently, the RNA samples were stored at −80° until processing. RNA-seq data were analyzed as described in Ouonkap et al. 2024^19^. All RNA-seq data are available on the Sequence Read Archive (SRA) at https://www.ncbi.nlm.nih.gov/sra under the accession number [PRJNA980666, PRJNA611819].

### Protein extraction and analysis

For protein extraction, pollen tubes were collected and the media was removed by centrifugation at 13000xg for 15 minutes. The cells were then ground using a TissueLyser machine. Subsequently, 200 μL of protein extraction buffer (5% SDS, 50mM Tris HCl, and 8M Urea, pH 7.55) was added. The mixture was incubated on ice for 30 minutes. To remove cell debris, the lysate was clarified by centrifugation at 13,000 x g for 10 minutes at 4°. The supernatant containing soluble proteins was carefully transferred to a tube on ice for analysis. The protein concentration was measured with the Bradford assay (Bio-Rad). The lysate was stored at –80° until processing.

Sample preparation for proteomic analysis using S-Trap method Samples were reduced using Dithiothreitol (DTT, 10mM final concentration, Sigma Aldrich, 646563-10X.5ML) and incubated at 55° for 45 minutes with mixing. Following incubation, samples were alkylated using Iodoacetamide (IAM, 20mM final concentration, Thermo Scientific, A39271) and incubated at room temperature in the dark for 30 minutes. Following alkylation, samples were acidified using Phosphoric Acid (2.5% final concentration, Sigma Aldrich, 79617-250ML), and vortexed for 30 seconds. 900mL of ice-cold Binding Buffer (9 parts LC/MS Grade Methanol (ACROS Organics, 61513-0025):1 part 1M TEAB (100mM Final concentration)) was added to each sample before the entire sample was transferred to each S-Trap spin column and centrifuged for 30 seconds at 4000xg. Each sample was then washed once with 150mL of binding buffer. Following protein washing, 1 vial (20mg) of Trypsin (Promega, V5111) was reconstituted in 50mM Triethylammonium Bicarbonate (TEAB, Sigma Aldrich, T7408) and added to each micro spin column to yield a final concentration of 1mg of trypsin for 20mg of protein in 100mL. Samples were then moved to a humidified chamber in a 37° incubator and left overnight (18hrs). On Day 2, samples were left to cool to room temperature for 15 minutes before 40mL of 50mM TEAB was added and each sample was centrifuged for 1 minute at 4000xg. Peptides were eluted with 40mL of LC/MS grade Water (Honeywell, 39253-4X2.5L) and 0.1% Formic Acid (Fisher Scientific, A117-50), and centrifuged again for 1 minute at 4000xg. Finally, to completely elute hydrophobic peptides, 40mL of 50% LC/MS Grade Acetonitrile (Fisher, A955-4) was added to each sample and allowed to stand for 5 minutes before being centrifuged for 1 minute at 4000xg. Each sample was concentrated using a speed vacuum concentrator (Thermo Fisher Scientific) for approximately 90 minutes. With solvent removed, samples were reconstituted in 100mL of Solvent A (Water with 0.1% Formic Acid) and spiked with indexed retention time peptides (Biognosys, Ki-3002-1) in a 1:50 dilution to monitor HPLC performance.

### LC-MS proteomic analysis

Samples were then injected onto a QExactive Orbitrap LC/MS system (Thermo Fisher Scientific) and separated over a 120-minute gradient consisting of solvent A (water + 0.1% Formic Acid), and solvent B (Acetonitrile and 0.1% Formic Acid) using a split-flow set-up on an Agilent 1200 series HPLC system. A 15cm long (75mm ID) capillary analytical column packed with XSelect CSH C18 2.5 mm resin (Waters) from the University of Arkansas for Medical Sciences (UAMS) IDeA National Resource for Quantitative Proteomics was used to separate digested peptides. The 120-minute gradient consisted of 95% Solvent A for 1 minute, 70% solvent A for 94 minutes, 5% solvent A for 6 minutes and finally 100% solvent A for the remaining 20 minutes all at a flow rate of 0.240 ml/min. The mass spectrometer acquisition utilized a Full MS/ddMS2 (Top9) centroid experiment. Full MS parameters used a default charge state of 2, 1 microscan, resolution of 70,000, AGC target of 3e6, 200ms Maximum Injection time, in a scan range of 400-1800m/z. ddMS2 parameters were 1 microscan, resolution of 17,500, AGC target of 2e4, a max injection time of 200ms, a loop count of 9, Top 9 precursors, Isolation window of 2.5m/z, collision energy of 28, and a scan range of 200-2000 m/z. Data dependent settings consisted of a minimum AGC target of 2e2, 1e3 intensity threshold, unassigned charge exclusion, all charge states, preferred peptide match, isotope exclusion, 30 seconds of dynamic exclusion.

### Label free quantitation and MaxQuant search parameters

Raw files were processed using MaxQuant (Ver 2.1.4) using 1% peptide and protein FDR search constraints, all values were default except for 20ppm first search tolerance and 7 ppm main search tolerance against Heinz Tomato protein fasta file (ITAG4.1). Variable modifications consisted of just oxidation (M) and protein N-terminal Acetylation. Carbamidomethylation was the sole static modification for the Trypsin/P search. Match between runs algorithm was unselected and intensity-based absolute quantification (iBAQ) was selected. The proteingroups.txt file output was further processed in MS Excel and Relative IBAQ was calculated as the intensity of each protein divided by the sum of all intensities for each given sample. In total, 3,028 proteins were detected in Heinz, 2,602 in Nagalang, and 2,741 in Tamaulipas, highlighting the high quality of the extracted material. The mass spectrometry proteomics data have been deposited to the ProteomeXchange Consortium via the PRIDE ^56^ partner repository with the dataset identifier **PXD053410**. DEqMS package was used to calculate differential protein expression (DEP) ^57^.

### Alignment and phylogenetic analysis of RALF peptides, receptors, and co-receptors

Protein sequences of RALF peptides were obtained from the Sol Genomics Network (ITAG4.1) for tomato and from the TAIR database (TAIR11) for Arabidopsis. Multiple sequence alignment (MSA) was performed using the MUSCLE method. The phylogenetic tree was constructed based on the neighbor-joining statistical method, with 100 bootstrap replicates. For the alignment of RALFs, predicted signal peptides and propeptides (if present) were removed, and only the predicted mature RALF sequences were used.

### Ralf peptide design

All peptides (Supplementary Table 3) used in this study were synthesized by Biomatik with purity >95%. They were diluted in ddH2O.

### Immunolocalization

Immunolabeling was performed as previously described by ^58^. Pollen grains were collected and allowed to germinate for 2 hours, after which pollen tubes were collected, fixed in paraformaldehyde-based fixation solution, and stored at 4° until use. Fixed pollen tubes were washed with PBS, incubated in a blocking solution, and then treated overnight at 4° with anti-LM19 or anti-LM20 antibodies for weakly or highly methylesterified HGs, respectively (Kerafast, https://www.kerafast.com). After washing, pollen tubes were incubated with Alexa488-conjugated secondary antibodies, followed by additional washes. Control samples were treated with the secondary antibody only. Immunolabeled pollen tubes were visualized using an inverted Nikon Ti2-E Widefield Fluorescence Microscope under identical imaging conditions, and fluorescence intensity was analyzed using ImageJ software.

### Generation of *RALF1* CRISPR/Cas9 Mutants

To generate CRISPR/Cas9-induced mutations in the *RALF1* gene, we used the CRISPR-P web tool (http://crispr.hzau.edu.cn/CRISPR2/) to design two guide RNAs. The guides were carefully selected to target regions upstream of the mature peptide or motif-encoding sequences, ensuring disruption of *RALF1* function. The CRISPR/Cas9 construct was assembled using the GoldenBraid cloning strategy. The final plasmid was then sent to Dr. Thomas Clemente’s laboratory at the University of Nebraska for Agrobacterium-mediated transformation. Seeds from Tamaulipas and Heinz cultivars were used for transformation to assess the effect of *RALF1* disruption in contrasting genetic backgrounds.

### Western Blot Analysis

Proteins were separated by SDS-PAGE on 4–12% acrylamide gels and transferred onto 0.22-μm nitrocellulose membranes ^59^. Running and transfer buffers were obtained from Invitrogen. Membranes were blocked and incubated overnight at 4° with rabbit polyclonal antibodies (Boster) against RALF1 (1:1000), and PEPC (1:3000) (ref) in 20 mM Tris-HCl (pH 7.4), 150 mM NaCl, 0.1% Tween-20, and 2% BSA. After washing, membranes were incubated for 1 h at room temperature with horseradish peroxidase–conjugated secondary antibodies. Protein expression was detected by chemiluminescence following the manufacturer’s instructions (ECL, Pierce 32106) and imaged using the ChemiDoc™ MP Imaging System (Bio-Rad Laboratories, Hercules, CA, USA). Images were analyzed with Image Lab. Software (Version 2.3.0.07).

---

**Supplementary Figure 1: Related to Figure 1.**
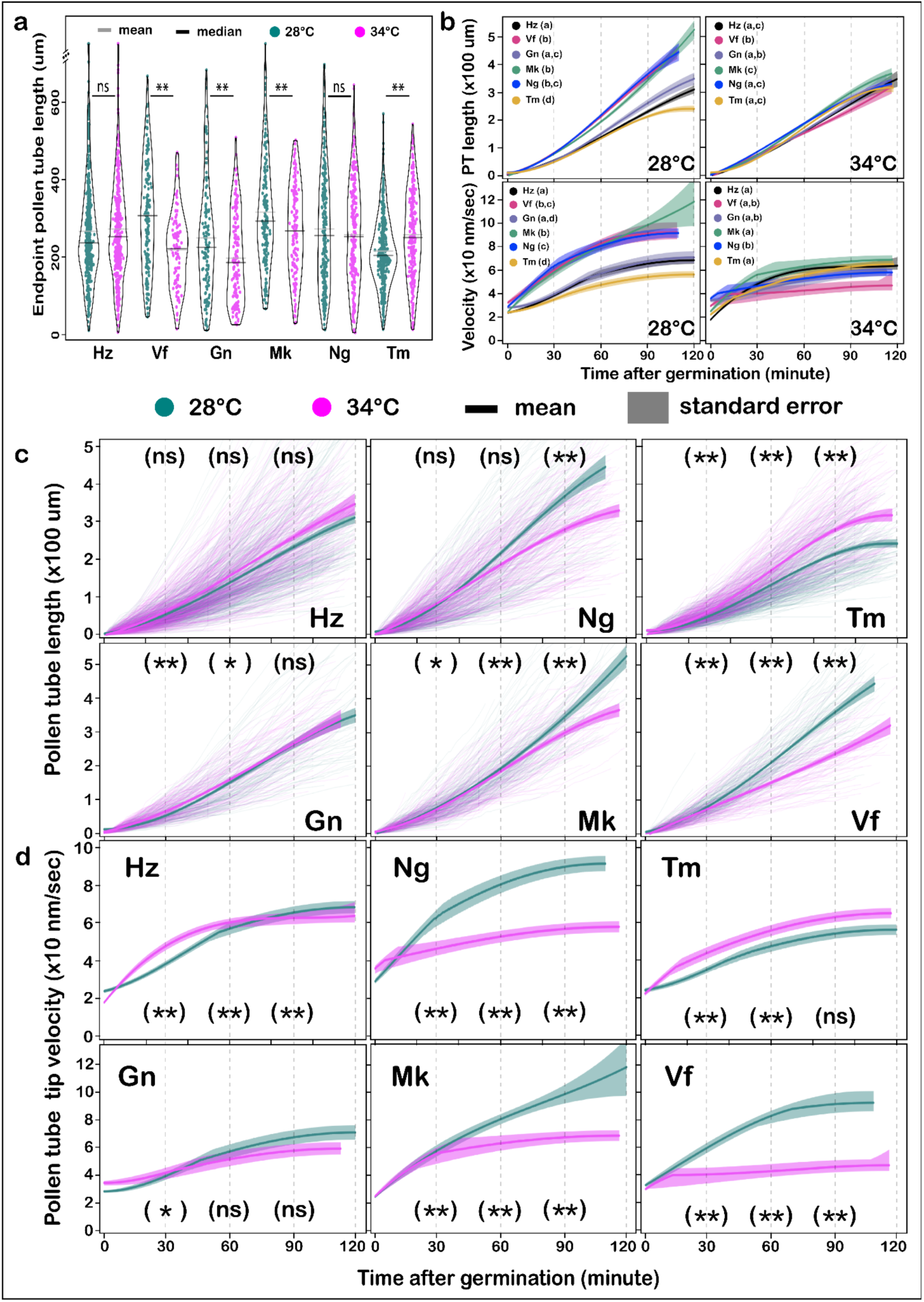
Effects of high temperature on pollen tube elongation and tip velocity. **(a)** Pollen tube length at 120 minutes without taking into account time of germination. Statistical analysis determined using the Welch Two Sample t-test with p.value ranges of ns>0.05>\**>0.01>*\*\*. **(b)** Comparison of the evolution of pollen tube length across all genotypes analyzed. Statistical significance was determined using the Welch Two Sample t-test at 90 minutes of growth. Shared letters indicate no statistical significance at p < 0.01. **(c)** Comparison of velocity across all genotypes analyzed. Statistical significance was determined using Dunn’s test. Shared letters indicate no statistical significance at p < 0.01. **(d)** Effects of temperature on pollen tube length over time. Statistical analysis was done at 30 minutes, 60 minutes, and 90 minutes after germination using the Welch Two Sample t-test using p.value ranges of ns>0.05>\**>0.01>*\*\*. **(e)** Effects of temperature on pollen tube velocity. Statistical analysis was performed as described in (d) but using the Mann-Whitney-Wilcoxon U test.

**Supplementary Figure 2: Related to Figure 1.**
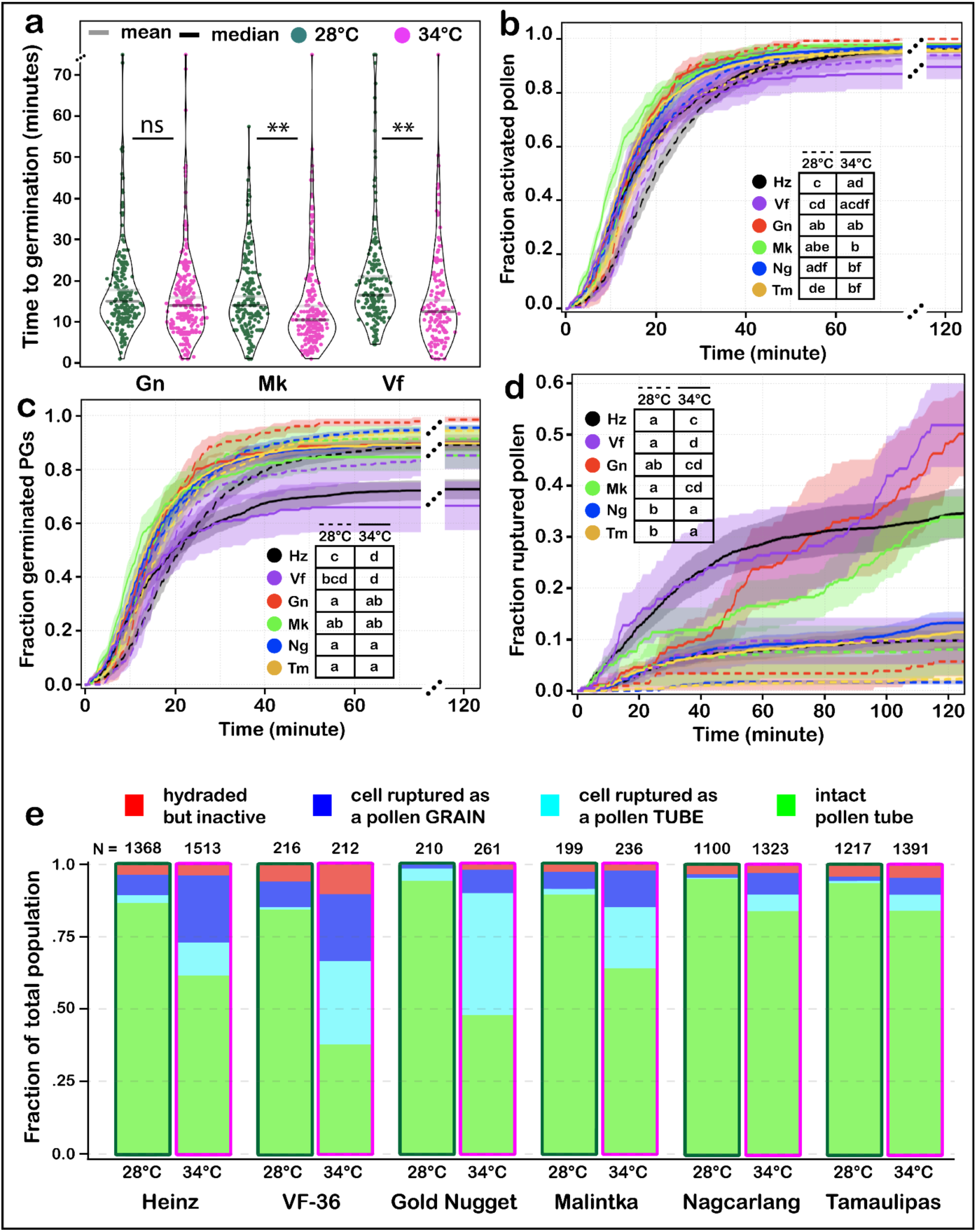
Effects of High temperature on pollen performance. **(a)** Time to successful germination at indicated temperatures for gold nugget (Gn), Malintka (Mk), and VF-36 (Vf). Each dot represents a germinated pollen grain. Statistical analysis was performed using the Mann-Whitney-Wilcoxon U test using p.value thresholds of ns > 0.01 > * > 0.001 > **. **(b)** Fraction of activated pollen grains over time. **(c)** Fraction of successfully germinated pollen grains over time. **(d)** Fraction of ruptured pollen grains and tubes over time. For **a-d,** Statistical analysis was performed using survival analysis (methods) with a time cutoff of 120 minutes and pairwise p.values were bonferroni adjusted. Similar letters in tables represent no statistical significance between groups at a p.value < 0.01. Different letters in tables represent statistically significant differences at a p.value < 0.01 between groups. **(e)** Cumulative barplots showing the density of each pollen performance population tracked.

**Supplementary Figure 3: Related to Figure 2.**
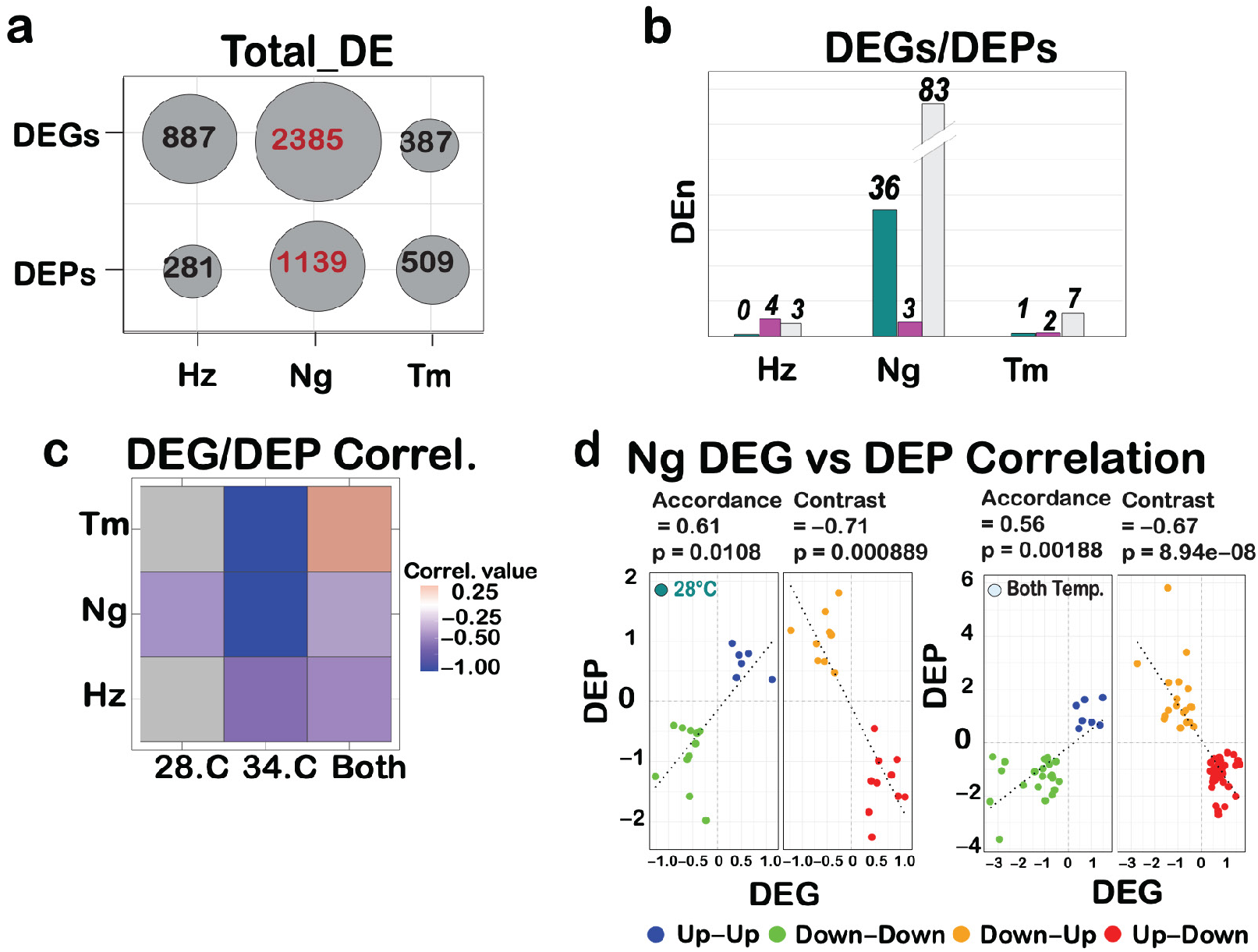
Negative Correlation Between Transcript and Protein Levels. (c) Heat map showing the correlation between DEPs and DEGs across three tomato cultivars under 28°, 34°, and both temperature conditions. Color intensity represents the Spearman correlation coefficient between transcriptomic and proteomic changes. A value of –1 indicates a strong negative correlation, while gray reflects insufficient overlap or undetectable correlation due to limited shared genes or proteins. (d) Transcript-protein correlation among shared DEGs and DEPs in Nagcarlang at 28° (n = 36) and both temperature conditions (n = 83). Each point represents a gene-protein pair, colored by the direction of transcript and protein regulation: Up-Up (blue), Down-Down (green), Up-Down (red), and Down-Up (orange). Genes with concordant expression (Up-Up and Down-Down) showed a moderate positive Spearman correlation (ρ = 0.56, p = 0.0018), while those with opposing regulation (Up-Down and Down-Up) exhibited a strong negative correlation (ρ = −0.71, p = 0.0001).

**Supplementary Figure 4: Related to Figure 2.**
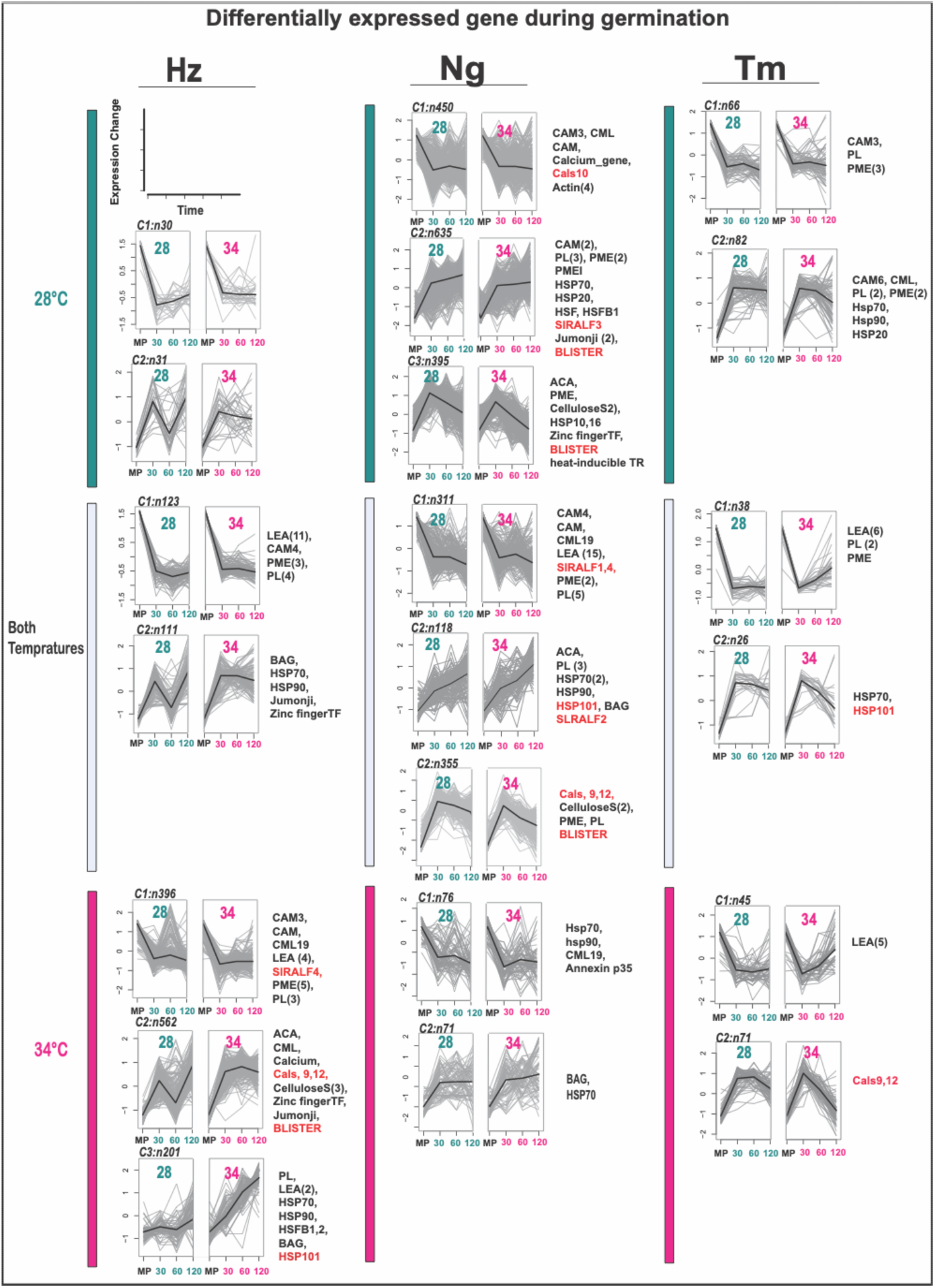
transcriptomic profile analysis of pollen germination in different tomato cultivars. Clustering analysis of DEGs identified in Figure 3, Panel C, representing the transcriptomic changes during pollen germination in various tomato cultivars. Each cluster reflects a distinct pattern of protein expression across the different varieties. Cellulose S, Cellulose Synthase; CAM, Calmodulin; CML, Calmodulin-Like protein; PME, Pectin Methylesterase; PL, Pectate Lyase; PG, Polygalacturonase; LEA, Late Embryogenesis Abundant protein; BAG, Bcl-2-Associated Athanogene family protein; Heat-induced TR, Heat-Induced Transcriptional Regulator; Blister, Blister-Like Protein.

**Supplementary Figure 5: Related to Figure 2.**
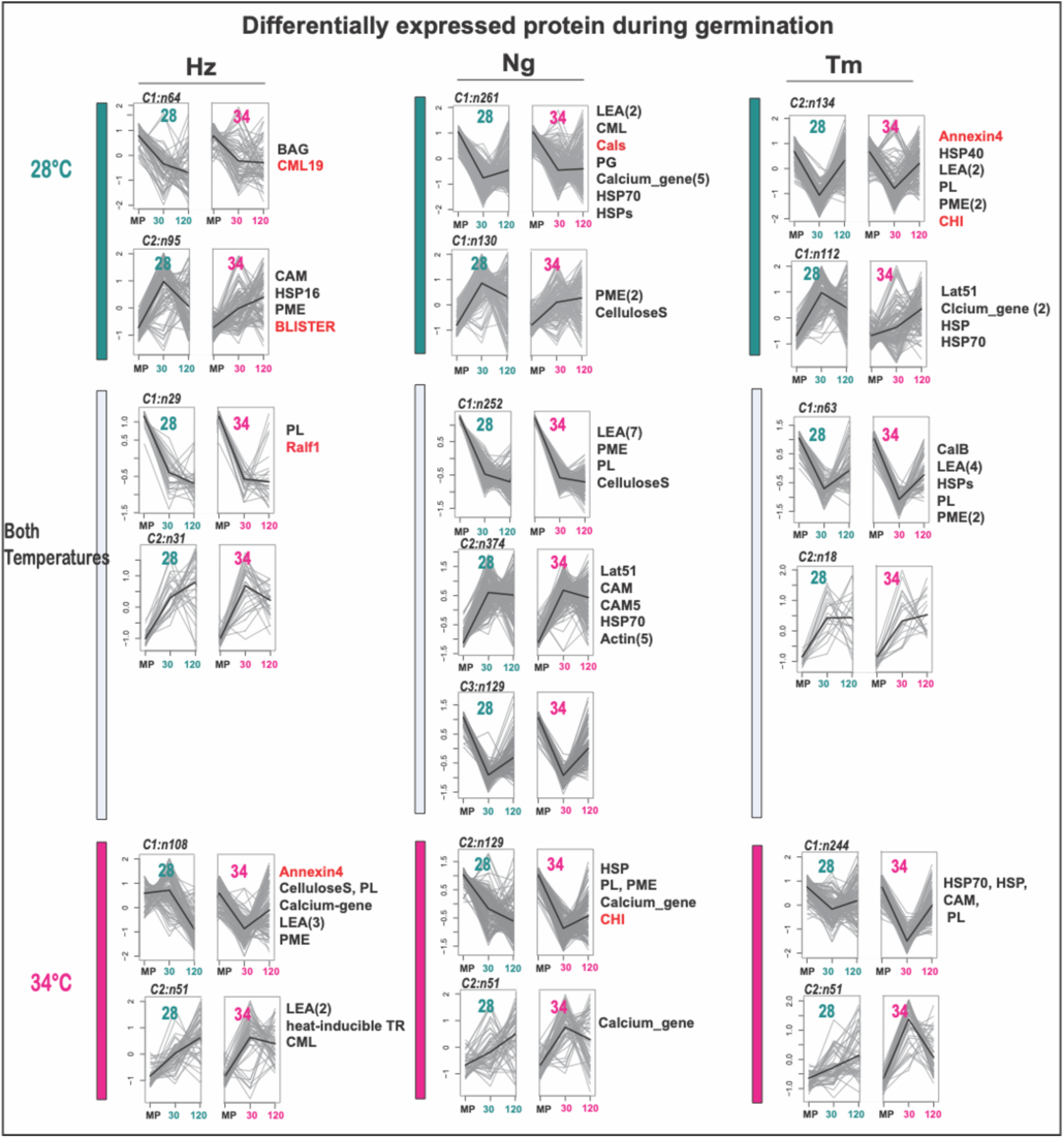
Proteomic profile Analysis of pollen germination in different tomato cultivars. Clustering analysis of DEPs identified in Figure 3, Panel C, representing the proteomic changes during pollen germination in various tomato cultivars. Each cluster reflects a distinct pattern of protein expression across the different varieties. Cellulose S, Cellulose Synthase; CAM, Calmodulin; CML, Calmodulin-Like protein; PME, Pectin Methylesterase; PL, Pectate Lyase; PG, Polygalacturonase; LEA, Late Embryogenesis Abundant protein; BAG, Bcl-2-Associated Athanogene family protein; Heat-induced TR, Heat-Induced Transcriptional Regulator; Blister, Blister-Like Protein.

**Supplementary Figure 6: Related to Figure 2.**
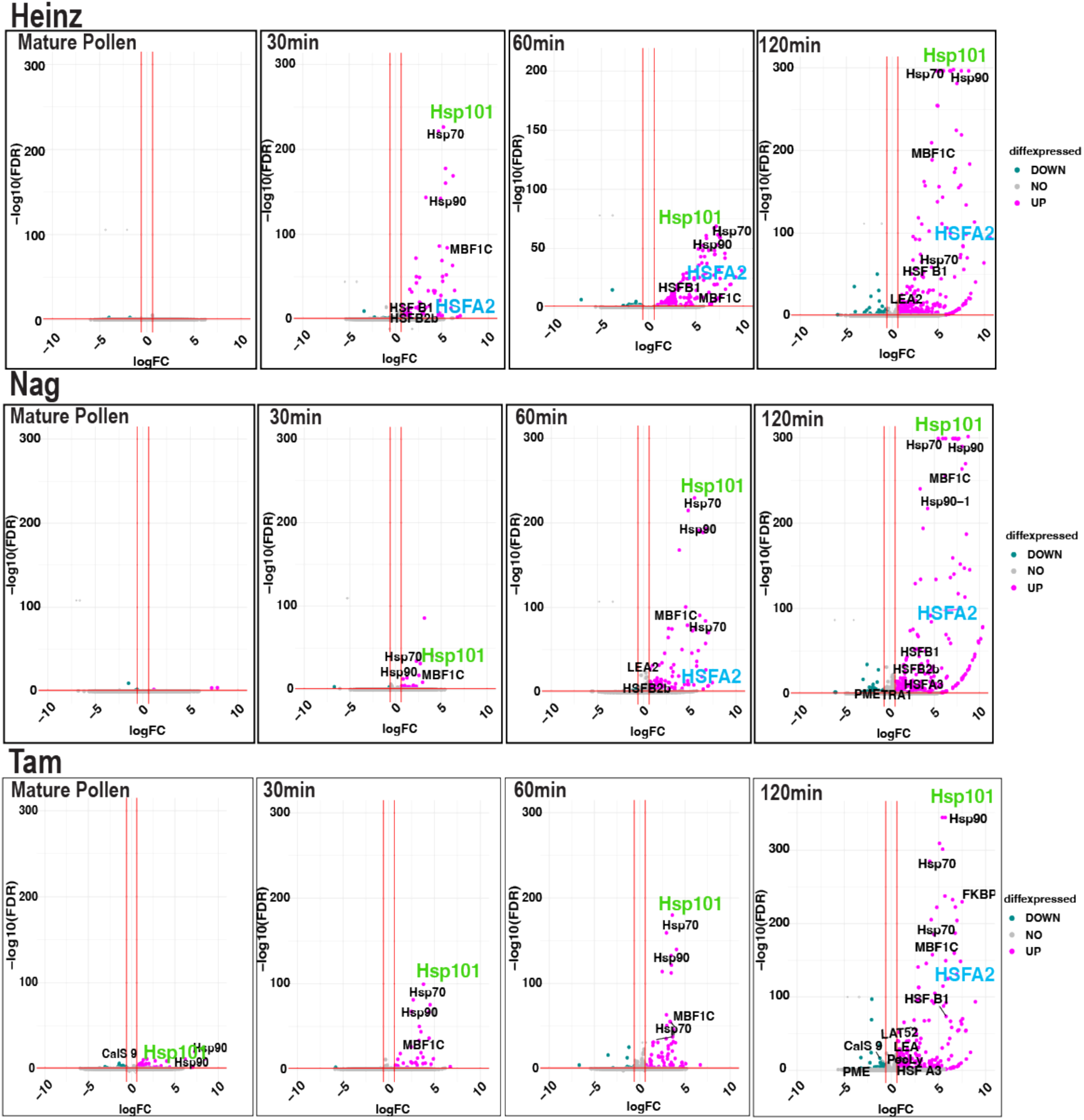
Volcano plot showing DEGs between 28° and 34° across cultivars at different time points. Genes with an absolute log fold change ≥ 0.5 and an adjusted p-value < 0.05 were considered significantly differentially expressed. Upregulated genes (logFC ≥ +0.5) are highlighted in magenta, and downregulated genes (logFC ≤ –0.5) are shown in cyan. Selected heat stress–related genes, including HSP101, HSP90, HSP70, HSFA2, HSFB1, and MBF1C are highlighted in the plot to indicate their dynamic expression in response to heat conditions.

**Supplementary Figure 7: Related to Figure 3.**
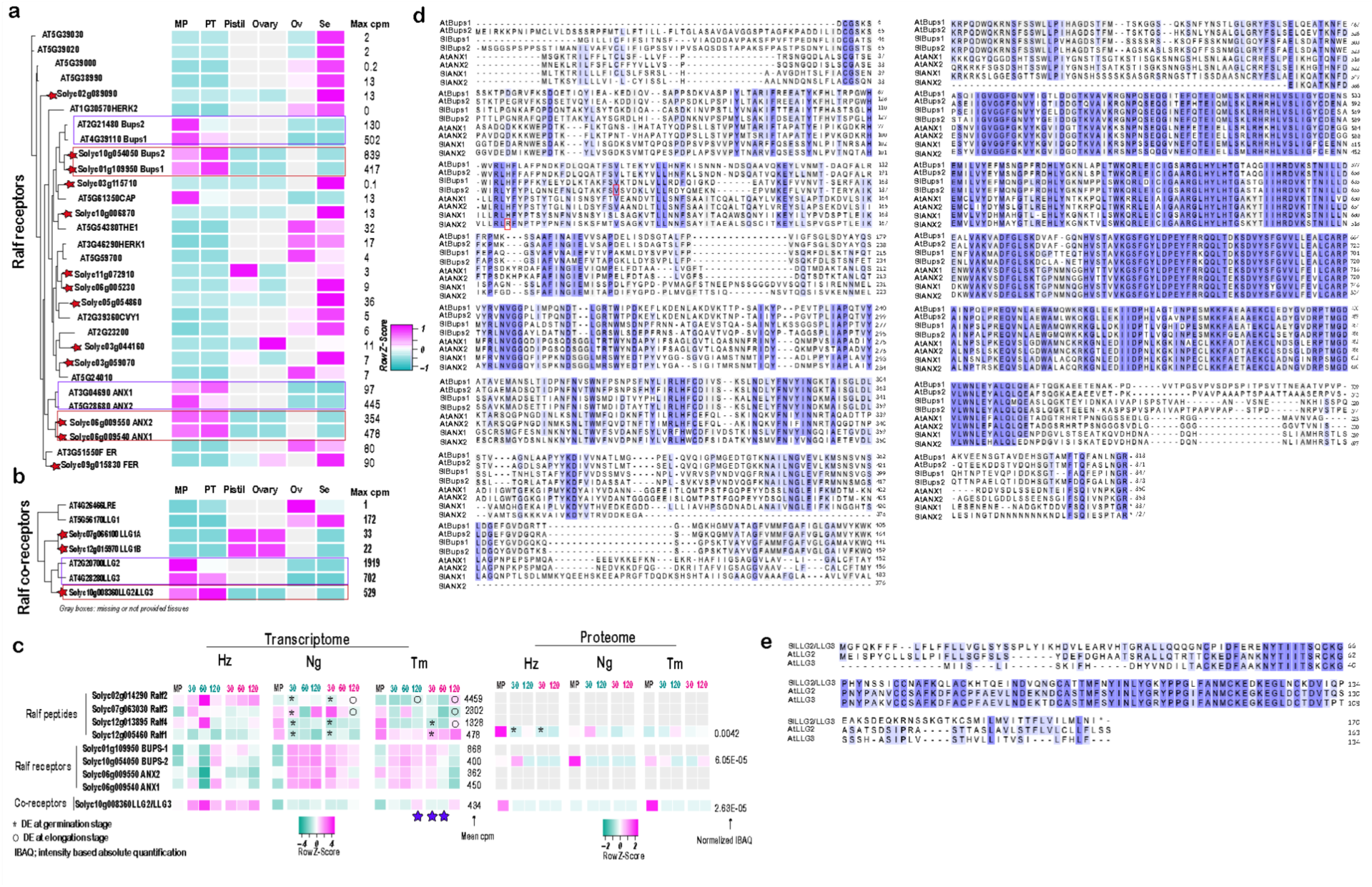
Tomato pollen expresses four RALF receptors and two co-receptors. The phylogenetic tree illustrates the evolutionary relationships between tomato and Arabidopsis Ralf receptors **(a)** and co-receptors **(b)**. Additionally, a heatmap shows the expression levels (RNA) of these genes in different tissues such as mature pollen "MP", pollen tube "PT", pistil, ovary, ovule "Ov" and seedling "Se". Max cpm represents the highest expression value observed across the different tissues. Gray boxes in the heatmap indicate tissues for which data are missing or not provided. **(d, e)** Multiple sequence alignments of the most abundantly expressed RALF receptors (c) and co-receptors (d) in mature pollen and the pollen tube, compared with their Arabidopsis homologs. Conserved amino acid residues are shaded. In panel (c), red boxes highlight genotype-specific amino acid variations observed in the Nagcarlang: arginine (R) residue in SlANX2 and a valine (V) in SlBUPS2 in the Hz correspond to a histidine (H) and an alanine (A), respectively, in the Ng cultivar.

**Supplementary Figure 8: Related to Figure 4.**
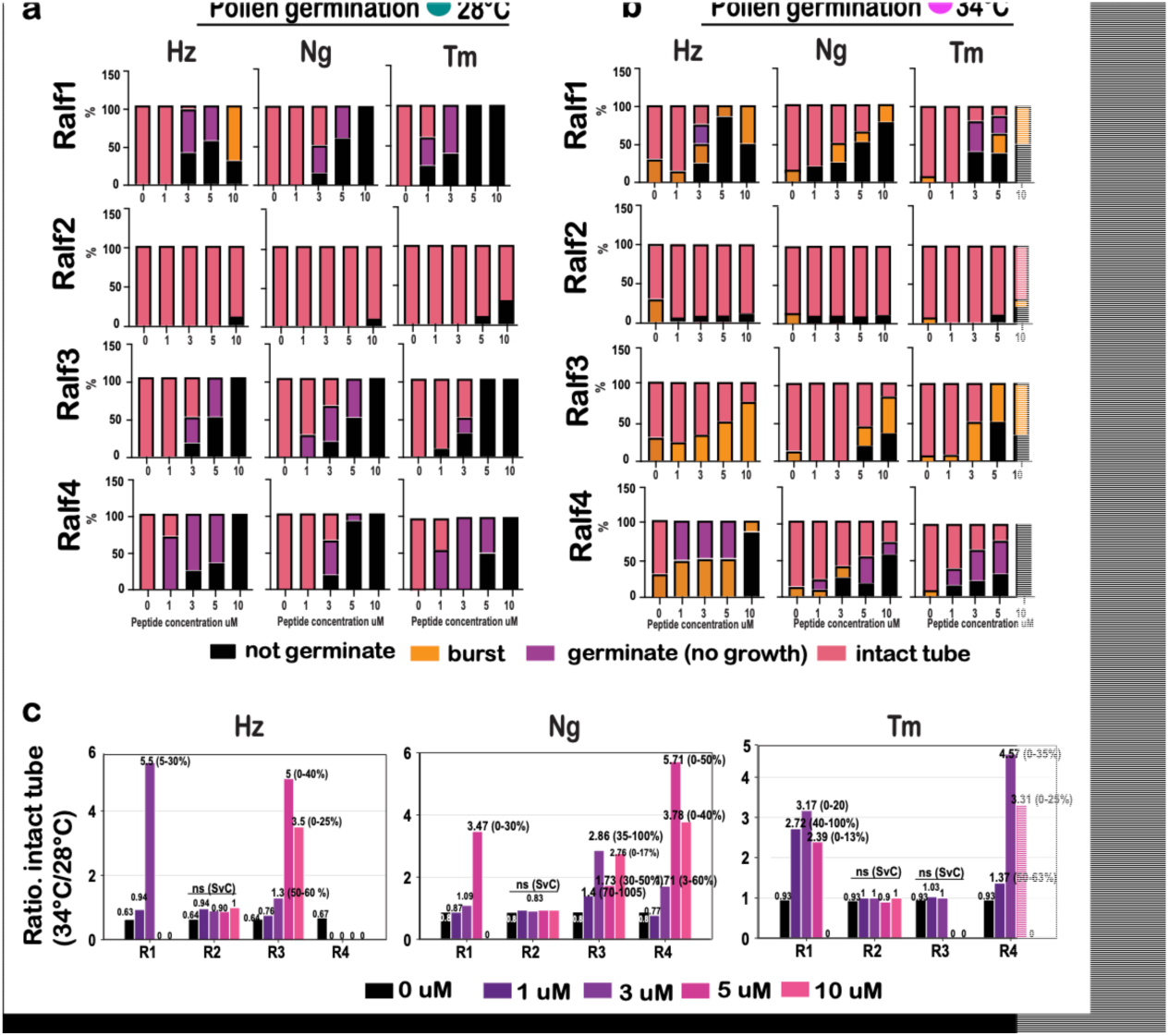
Effects of different RALF peptides on pollen tube germination under **(a)** 28° or 34°**. (c)** Ratio of percentage of pollen tube formation between control and heat stress, a ratio greater than 1 indicates an enhancement of pollen tube formation under heat stress compared to control (%) represents the increase of pollen tube formation by HT. We grew Heinz, Nagcarlang, and Tamaulipas pollen with each peptide at 1, 3, 5, and 10 μM for 90 minutes at 28° and evaluated pollen germination and tube elongation. 10 uM SlRALF4 and SlRALF3 completely inhibited pollen germination of all three cultivars, whereas 10 uM SlRALF1 inhibited germination in Nagcarlang and Tamaulipas, but caused ∼80% of pollen to rupture in Heinz. SlRALF1, 3, and 4 all showed dose-dependent inhibition of pollen germination between concentrations of 1 and 10 uM that varied across the three cultivars. SlRALF2 had little effect on pollen germination even at 10 uM. These data suggest that SlRALFs 1, 3, and 4, but not SlRALF2 control pollen germination and that responses vary across cultivars. These findings are consistent with previous analysis of SlRALF3 (SlPRALF^24^). To test whether RALF signaling interacts with temperature, we conducted these experiments at HT (34°). Unlike our observations at 28°, 10 uM SlRALF3 did not completely inhibit pollen germination of any of the cultivars at 34°. Instead, at HT significant numbers of pollen burst (Heinz, Nagcarlang, and Tamaulipas) or produced an intact pollen tube (Heinz and Nagcarlang). The germination-blocking effects of SlRALF1 and SlRALF4 were also partially reversed at 34°. Again, SlRALF2 treatment had little effect on pollen tube growth at 34°. These data suggest that HT is antagonistic to RALF (SlRALF1, 3, and 4) signaling.

**Supplementary Figure 9: Related to Figure 4.**
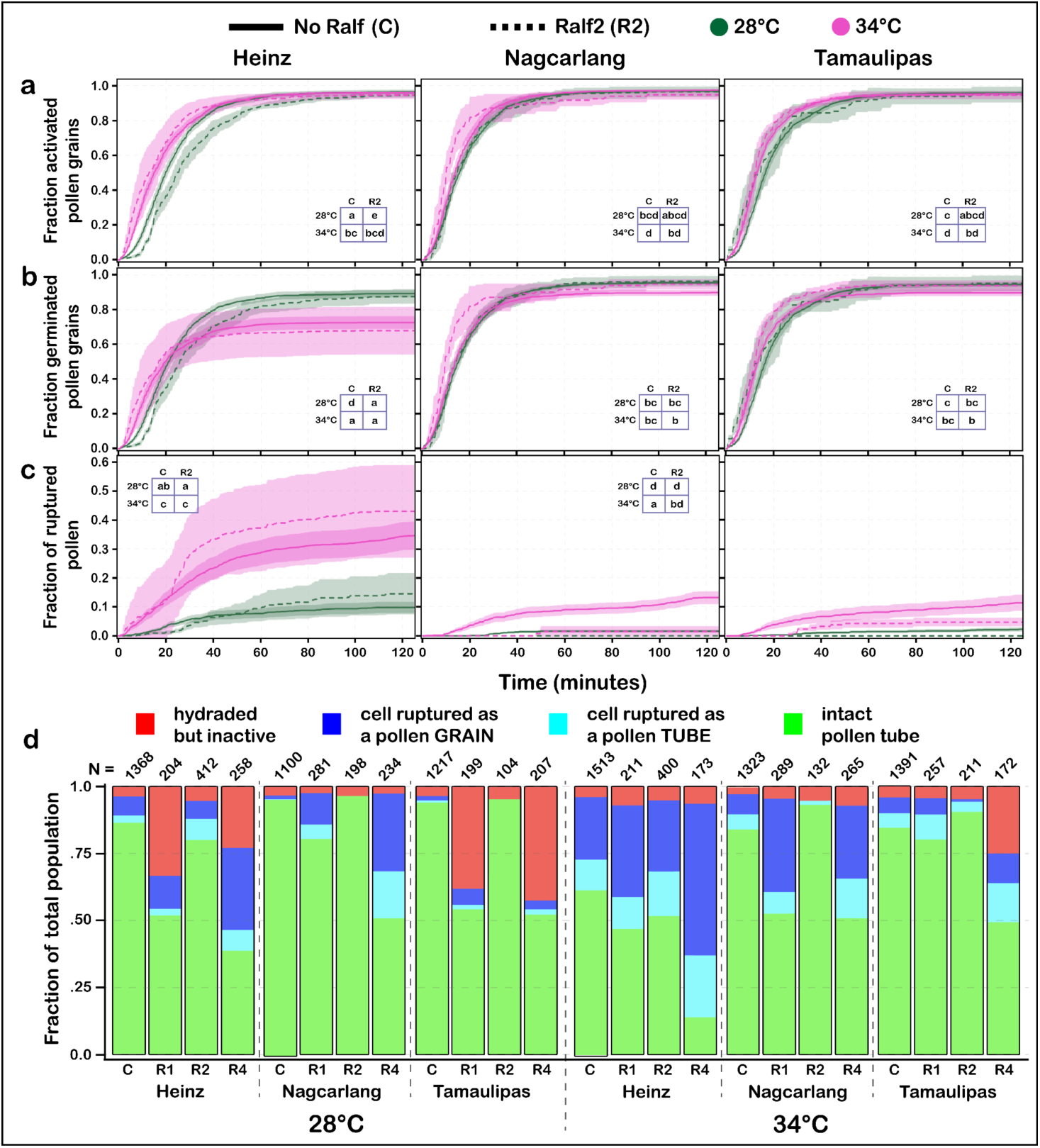
Effects of SlRalf2 on pollen grain activation, germination, and cell rupture. **(a)** Fraction of activated pollen grains over time – grains that exited the hydrated state (Figure 1a, red). **(b)** Fraction of successfully germinated pollen grains over time (Figure 1a, red to green). **(c)** Fraction of ruptured pollen grains and tubes over time (Figure 1a, blue + cyan). For c-e, Statistical analysis was performed using survival analysis (methods) with a time cutoff of 125 minutes and pairwise p.values were bonferroni adjusted. Similar letters in tables represent no statistical significance between groups at a p.value < 0.01. Different letters in tables represent statistically significant differences at a p.value < 0.01 between groups. **(d)** Cumulative barplots showing the density of each pollen performance population tracked.

**Supplementary Figure 10, related to Figure 6:**
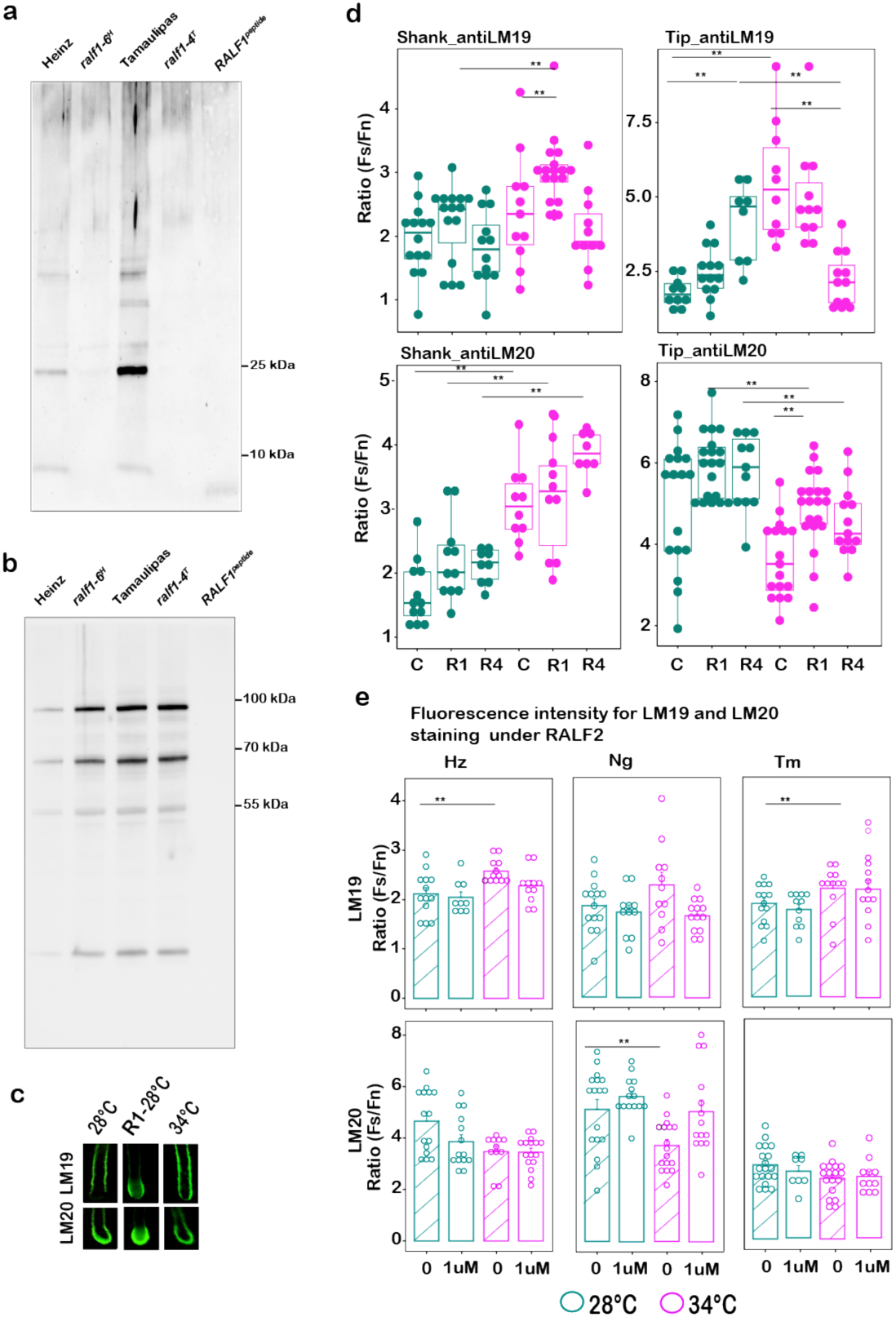
**(a, b)** Complete Western blot analysis using anti-RALF1 and anti-PEPC antibodies, respectively, in WT Heinz, WT Tamaulipas, mutant lines, and synthetic RALF1 peptide. PEPC was used as a loading control. **(c)** Representative images of LM19 and LM20 immunostaining, showing pectin distribution in the shank and tip regions of pollen tubes in Nagcarlang. RALF1 (R1) treatment is used as a representative for staining patterns. **(d)** Quantification of fluorescence intensity for LM19 and LM20 staining in the shank and tip region under RALF1 and RALF4 treatments in Nagcarlang, and at 23° and 34°. Fluorescence intensity was normalized using the ratio of LM19 or LM20-stained fluorescence (Fs) to fluorescence of unstained controls (Fn). Statistical significance was determined using Dunn test and indicated as p < 0.01 (**), only. **(e**) Quantification of fluorescence intensity for LM19 staining in the shank region and LM20 staining in the tip under RALF2 treatment across cultivars at 23° and 34°. Fluorescence intensity was normalized using the ratio of LM19- or LM20-stained fluorescence (Fs) to the fluorescence of unstained controls (Fn). Statistical significance was determined using the Dunn test and indicated as **p < 0.01. Only significant values are shown in the graph (indicated by two asterisks).

## Abbreviations

HT: high temperature
Hz: Heinz
Mk: Malintka
Ng: Nagcarlang
Tm: Tamaulipas
RALF: Rapid Alkalinization Factor

## Acknowledgements

This project was funded by grants from the US NSF (IOS-1939255) and the USDA/NIFA (2020-67013-30907). We thank Sherry Warner and Nick Vasquez (Brown University) for expertise in growing and maintaining tomato plants.

## Author Contributions

R.A.A., S.V.Y.O., J.I., Y.O., M.A.A., C.T., A.H., N.O., G.W., A.D. J.P., M.M., N.D., and M.A.J. conducted the experiments and developed the methods. R.A.A., S.V.Y.O., and M.A.J. analyzed the data. R.A.A., S.V.Y.O., and M.A.J. designed the experiments. R.A.A., S.V.Y.O., and M.A.J. wrote the manuscript.

